# Higher aboveground carbon stocks in mixed-species planted forests than monocultures – a meta-analysis

**DOI:** 10.1101/2022.01.17.476441

**Authors:** Emily Warner, Susan C. Cook-Patton, Owen T. Lewis, Nick Brown, Julia Koricheva, Nico Eisenhauer, Olga Ferlian, Dominique Gravel, Jefferson S. Hall, Hervé Jactel, Carolina Mayoral, Céline Meredieu, Christian Messier, Alain Paquette, William C. Parker, Catherine Potvin, Peter B. Reich, Andy Hector

## Abstract

Natural forest is declining globally as the area of planted forest increases. Planted forests are often monocultures, despite results suggesting that higher species richness improves ecosystem functioning and stability. To test if this is generally the case, we performed a meta-analysis of available results. We assessed aboveground carbon stocks in mixed-species planted forests vs (a) the average of constituent species monocultures, (b) the best constituent species monoculture, and (c) commercial species monocultures. We investigated whether any advantage of mixtures over monocultures was positively related to species richness, as well as potential mechanisms driving differences in carbon stocks between mixtures and monocultures. The meta-analysis dataset included 79 comparisons from 21 sites. Carbon stocks in mixed planted forests were higher than the average of stocks in monocultures of their constituent species, containing on average 70% more carbon. Mixed planted forests also out-performed commercial monocultures, containing on average 77% more carbon. There was c.25% more carbon in mixed planted forests relative to the best performing monocultures, although this difference was not statistically significant. Overyielding was highest in four-species mixtures (richness range 2-6 species). More data providing better coverage of richness and age gradients (study sites aged 3.5-28 years) is needed to increase confidence in these results. None of the potential mechanisms we examined (nitrogen-fixer present vs absent; native vs non-native/mixed origin; tree diversity experiment vs forestry plantation) consistently explained variation in the diversity effects. This suggests that our findings are driven by a combination of small (statistically insignificant) effects from these sources or further unidentified mechanisms or some combination of the two. We conclude that increasing tree species richness in planted forests can increase carbon stocks while bringing other potential benefits associated with diversification. However, implementation will depend on the balance of these benefits relative to the operational challenges and costs of diversification.

## Introduction

Globally, natural forest extent is declining, but the area of planted forest is increasing and currently comprises 7% of global forest cover (FAO, 2020). Planted forests comprise both plantation forests, which usually contain one or two species and are managed intensively for production purposes, and other planted forests, containing one or more tree species and managed less intensively for multiple uses (Messier et al., 2021). There is growing momentum to restore forest cover for climate change mitigation, biodiversity conservation, and sustainable development goals (Di Sacco et al., 2021; Griscom et al., 2017; Seddon et al., 2020), but many national forest restoration commitments focus on monoculture plantations. Indeed, 45% of forest restoration commitments from tropical countries involve establishing monoculture plantations, despite diversification of plantations being a key policy recommendation in a recent IPBES-IPCC report (Lewis et al., 2019; Pörtner et al., 2021). Experimental and theoretical work demonstrates that diverse plant systems are more productive and stable through time compared to monocultures, as well as better able to support diverse animal assemblages and provide other critical ecosystem services (Cardinale et al., 2012; Messier et al., 2021; Tilman et al., 2014). With the increasing calls for diversification of planted forests and the increasing number of studies addressing the effect of planted tree diversity on productivity, it is timely to assess whether carbon stocks in mixed-species planted forests can equal or exceed monocultures (Messier et al., 2021). Without this evidence, forestry practitioners lack the certainty needed to deviate from the more commonly deployed and understood monoculture systems.

The global forest carbon stock is estimated at 662 Gt of carbon, with 295 Gt stored in living organic matter, 300 Gt in the soil and 68 Gt in dead wood and litter (FAO, 2020). At their maximum pre-agricultural extent global forests are estimated to have contained roughly 500 Gt of carbon in living biomass and 700 Gt of carbon the soil (Malhi et al., 2002). Restoration of forest cover could recover large amounts of this carbon, with global reforestation efforts capable of sequestering c.10 GtCO_2_e a year (Griscom et al., 2017). However, the method of reforestation used has a strong influence on the carbon sequestered within the forest, with monoculture plantations being more limited in their potential to store carbon (Lewis et al., 2019). Future planted forests can contribute to climate change mitigation through terrestrial carbon storage and the substitution of wood products for fossil fuels and high-carbon products such as concrete (Forster et al., 2021). New planted forests must be optimised for carbon storage and to maximise other benefits, and diversification could be one route to achieving this.

Multiple studies have demonstrated a positive relationship between diversity and productivity in natural (e.g. Liang et al., 2016; Xu et al., 2020) and planted forest systems (e.g. Ewel et al., 2015; Huang et al., 2018). Meta-analyses have shown the existence of overyielding in mixed forests, with primary production exceeding the average productivity of component monocultures by about 15-20% (Jactel et al., 2018; Zhang et al., 2012). Our work builds on these previous meta-analyses, which assess the effect of tree diversity on carbon accumulation in planted and naturally occurring forests (Zhang et al., 2012), biodiversity-productivity relationships in forests and their relation to climate (Jactel et al., 2018) and growth rates in mixed vs. monoculture plantations (Piotto, 2008). By focussing solely on planted forests and assessing carbon stocks, we address this question in the context of the increasing interest in tree planting for carbon sequestration.

Forests established with higher numbers of tree species would therefore be expected to store more carbon in aboveground biomass than less diverse forests (Jucker et al., 2014). Moreover, although monoculture species are often selected for their high yield and ease of management (Nabuurs et al., 2018), they may lack resilience to perturbation (Jactel et al., 2017, 2021), which could compromise their long-term carbon storage potential (Hutchison et al., 2018; Messier et al., 2021; Osuri et al., 2020), especially under future climate conditions. For example, in a tree planting experiment over 15 years in Panama, growth in mixed stands was more stable in response to climatic extremes, and tree mortality was lower, compared to monocultures (Hutchison et al., 2018). There is also evidence that carbon capture can be more stable through time and recover more rapidly after drought in species-rich natural forests, compared to species-poor plantations (Osuri et al., 2020; Pardos et al., 2021). The simple act of mixing species with different functional traits can also result in more reliable levels of establishment, compared to a mosaic of monocultures of different species, where some may perform poorly (Tuck et al., 2016). Finally, diversification of forests is expected to increase resistance to pests and disease (Gamfeldt et al., 2013; Jactel et al., 2021; Jactel & Brockerhoff, 2007; van der Plas et al., 2016), which could also improve carbon stability and storage.

Alongside increased resilience to perturbation, there are other mechanisms that may lead to greater carbon accumulation in mixed-species planted forests compared to monocultures. Species mixtures may demonstrate “complementarity”, whereby niche differentiation or facilitation among individual species enhances overall performance (Loreau & Hector, 2001). For example, variation in crown architecture complementarity can achieve overyielding in mixed-species planted forests (Jucker et al., 2015; Williams et al., 2017). Complementarity may also be higher among native species that have coevolved (Cook-Patton & Agrawal, 2014; Zuppinger-Dingley et al., 2014). However, non-native plantation species might be expected to outperform native plantation species; plantation species (e.g., *Eucalyptus* and *Pinus* species) are often planted outside their native range and selected for their fast growth rates (Heryati et al., 2011; Marron & Epron, 2019). Species with particular traits can also enhance the overall productivity of a mixture. For example the presence of nitrogen-fixing species can enhance nitrogen availability for species unable to fix nitrogen, boosting carbon accumulation potential (Loreau & Hector, 2001; Mayoral et al., 2017). However, species mixing could lead to lower productivity in diverse stands where a lower yielding species dilutes a monoculture of a fast-growing, commercially valuable species. Mixing lower yielding species with a very high yielding species could lead to lower stand-level carbon accumulation because the diverse stand would fail to demonstrate “transgressive overyielding”, where diverse systems outperform even the most productive monoculture.

Planted forests can provide timber and other forest products, support local communities and provide an economic income, while helping to alleviate pressure on primary and semi-natural forest (McEwan et al., 2020; Messier et al., 2021). Given the dominant practice of establishing plantations as monocultures and the anticipated benefits of mixed-species planting for carbon accumulation (Beugnon et al., 2021) and biodiversity (Ampoorter et al., 2020), we need a robust examination of how carbon stocks compare in mixed planted forests relative to monocultures. Here, we present results of a meta-analysis, combining data from the Tree Diversity Network (TreeDivNet) (Verheyen et al., 2016) with data from peer-reviewed publications identified via a comprehensive literature search, to assess the effect of diversification on carbon stocks in planted forests, addressing the following research questions:

1. Are carbon stocks in mixed-species planted forests higher than in monocultures? We compared carbon stocks in mixed planted forests to carbon stocks in (a) the average of monocultures, (b) the most productive monoculture, and (c) monocultures of commercial timber species, within an experiment.
2. Is any advantage of mixtures over monocultures positively related to species richness in the mixtures?
3. What are potential mechanisms driving differences in carbon stocks between mixtures and monocultures? Specifically, we compared how responses changed in stands with and without nitrogen fixers and in native versus non-native stands. We also compared tree diversity experiments to forestry plantations, hypothesizing that experiments may better control confounding factors and thus may be more likely to demonstrate any positive effect of diversity.

## Methods

### Data collation

This meta-analysis used studies compiled in a previous systematic literature review. Cook-Patton et al. (2020) compiled 10,937 studies using a systematic keyword search in Web of Science (19 April 2017) for studies published since 1975: TOPIC: (biomass OR carbon OR agb OR recover* OR accumulat*) AND (forest) AND (restorat* OR reforest* OR afforest* OR plantation* OR agroforest* OR secondary*). This initial literature set was augmented to 11,360 studies by including additional papers referenced within the original studies and datasets from Oak Ridge National Laboratory, International Centre for Research in Agroforestry, and the Chinese Academy of Forestry. Abstracts were reviewed to identify studies describing reestablishment of forest cover; these were reduced to 1,400 studies which quantified carbon or biomass stocks. Of these, 62 studies compared carbon in mixed and monoculture planted forests.

We selected the subset of studies that met our criteria for inclusion: mixed and monoculture planted forests of the same age and containing a measure of aboveground carbon or biomass, monocultures composed of a constituent species of the mixed treatment, and a specific geographic location (to classify species as native or non-native and to classify commercial species). Studies also needed to contain necessary information (mean, standard deviation, sample size) to calculate the effect size for the meta-analysis, the standardised mean difference (SMD) or “Hedges’ g” (Hedges, 1981). A total of 9 of the 62 studies met these criteria (Table S1). Common reasons for exclusion were confounding variables in the mixed and monoculture treatments (e.g. mixed and monoculture stands of different ages) or failure to provide the metrics needed to conduct a meta-analysis. We included two additional studies from a meta-analysis by Zhang et al. (2012). We then augmented this literature-based dataset with seven additional datasets from TreeDivNet (https://treedivnet.ugent.be/), made available by TreeDivNet projects in response to a call for data, bringing the total dataset to 18 studies. Some studies included multiple experiments, producing 21 independent experiments from 18 studies. The studies often included multiple distinct mixed treatments (multiple species combinations or ratios between species), yielding a total of 79 monoculture-to-mixed comparisons. Each unique species combination (based on species composition or species ratios) was defined as a treatment. We extracted from each experiment: (1) mean aboveground carbon or biomass for each treatment (calculated in the original studies using allometric equations); (2) standard deviation of the carbon or biomass measure (in some cases calculated from standard error and sample size); (3) sample size for each treatment; (4) species composition of each treatment; (5) stand age and (6) geographic location. When data were presented graphically in the papers, we extracted estimates using WebPlotDigitizer (https://apps.automeris.io/wpd/).

### Data preparation

For research question 1a, where we were comparing carbon stocks in mixed planted forests to the average of carbon stocks in constituent species monocultures, this required us to calculate the mean and standard deviation across the monocultures relevant to each mixed treatment. In 9 of the 79 comparisons, only one of the mixed treatment constituent species was planted as a monoculture, so in these cases this monoculture was used in the comparison to the mixed treatment. Where two or more monoculture treatments were available, we calculated the average carbon/biomass and standard deviation. For the TreeDivNet data we had access to the full raw data, which was used to calculate the mean and standard deviation. For the studies resulting from the literature search, we used the extracted means and standard deviations for each monoculture treatment, to calculate a mean, standard deviation and sample size across monoculture treatments. The standard deviation was calculated based on the formula 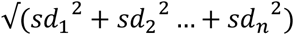 (Frey et al., 2006). The sample size of the average carbon/biomass measure was the aggregate sample size of all contributing monoculture treatments. The resulting means, standard deviations and sample sizes were used to calculate the SMDs.

For research questions 1b and 1c, the calculated (TreeDivNet studies) or extracted (studies from literature search) means, standard deviations and sample sizes from the mixed and monoculture treatments within each comparison were used to calculate SMDs. Question 1b compared carbon stocks in mixtures to the associated monoculture with the highest carbon/biomass stocks (n = 79 comparisons), again including the 9 comparisons where only a single monoculture was available. For research question 1c, we compared carbon stocks in mixtures to monocultures composed of commonly used commercial plantation species (n = 38 comparisons). We classified commercial species monocultures when there was evidence that these species were commonly used as commercial species in the study region (Table S2). We hypothesised that, as commercially grown tree species are often selected for their high yields, they may outperform the mixed treatments. The calculated (TreeDivNet studies) or extracted (studies from literature search) means, standard deviations and sample sizes were used to calculate SMDs for the comparison to the best or commercial species monocultures.

To assess some potential mechanisms behind differences in carbon stocks between mixed and monoculture treatments we classified each mixed treatment based on its species richness, presence of a nitrogen-fixing species in the mixture (yes/no), origin of the species (all native vs. some/all species non-native) and study design (designed experiment vs. existing forestry plantations).

To also present our results in real terms we calculated the percentage of monoculture carbon in the mixed treatment for each comparison, as SMD is on the scale of standard deviations. The carbon stocks in each treatment are presented in Tables S3 and S4. Where stocks were given as biomass in the original study, we converted these to carbon using the conversion factor of 0.47 (Aalde et al., 2006).

### Meta-analysis

All data manipulation, calculations and analysis were carried out in R (R Core Team, 2020). We used the *escalc* function in the package *metafor* to calculate the SMD (Hedges’ g) in carbon or biomass for each monoculture-to-mixed comparison and the associated sampling variance (Viechtbauer, 2010). Although some studies reported biomass stocks and others carbon stocks, we assume that carbon is a constant proportion of biomass (Aalde et al., 2006). Given this assumption, the calculation of the SMD therefore converts biomass and carbon onto the same scale. We hereafter refer solely to aboveground carbon stocks in the Results and Discussion. We calculated SMDs of the mixture treatment relative to the average of the monoculture treatments (overyielding), the monoculture treatment with the highest carbon stocks in a study (transgressive overyielding) and commercial monoculture treatments (when available).

Multi-level random-effects models with restricted maximum likelihood estimation were fitted using the function *rma.mv* in the *metafor* package (Viechtbauer, 2010). Many of our studies contained multiple mixed treatments which were compared to the same monoculture controls (71 out of 79 comparisons shared a monoculture with at least one other comparison). We included a random effect for study site to account for this. For the comparison to the best monoculture, we also included a random effect for monoculture treatment nested within study site. For the comparison to commercial monocultures, in some cases the same mixed treatment was compared to different monocultures and there were also cases where different mixed treatments were compared to the same monoculture; we included a random effect grouping by either shared mixed or monoculture treatment, nested within study site. All comparisons within a study site were between planted forests of the same age, however study sites covered the age range 3.5 to 28 years, the random effect for study site therefore also accounts for this. A random effect for each comparison was also included, to allow heterogeneity within study sites. We assessed the significance of the SMDs by determining whether the 95% confidence intervals included zero.

To assess the effect of the level of diversity on SMD, we fitted a mixed-effects model with a moderator for species richness (2-6 species), as a discrete variable, and separately compared mixed vs. average of monoculture and mixed vs. best monoculture. For mixed vs. commercial species monoculture comparisons, there were insufficient data across the levels of species richness to fit it as a moderator. For the mixed vs. average of monocultures and mixed vs. best performing monoculture analyses we fitted separate models for two, four, and six species mixtures, to estimate the effect size for each level of species richness; mixtures with three and five species only had one monoculture-to-mixed comparison each. We also fitted a model to estimate the effect size for two species mixtures compared to commercial monocultures.

To test potential predictors of differences in carbon stocks, we fitted separate mixed-effects models with moderators for presence vs. absence of nitrogen-fixer in the mixed treatment, native vs. some or all non-native species in the mixed treatment and designed experiments vs. existing plantations. For these models we only used data from two-species mixtures, as this was the only species richness level with good representation of both factor levels (Fig. S1). To estimate the overall effect size for each moderator level, we subset the data by each level of the moderator and fitted separate models for: nitrogen-fixer present, nitrogen-fixer absent, native species, non-native/mixed origin species, designed experiments, and existing plantations.

Age may mediate the effect of diversity on carbon stocks, to assess this we fitted a mixed-effects model with a continuous moderator for stand age for the mixed vs. average, best and commercial monoculture comparisons. These models were fitted using data from two-species mixtures, as this was the only species richness level with good representation across the age range (Fig. S1a). A linear and quadratic fit were compared using AIC; the quadratic relationship was fitted post-hoc, based on visual inspection of the data.

We checked for publication bias using funnel plots fitted for each analysis (Supplement S2) and explored the removal of the most extreme positive and negative SMDs. The impact on the interpretation of our results is discussed in Supplement S2.

## Results

From a systematic review of over 11,360 publications and compilation of data from a global network of tree diversity experiments, we compiled a maximum dataset of 79 monoculture to mixed comparisons, from 21 sites. Within the 79 comparisons there were 51 unique mixed-species combinations from a pool of 54 species (Table S3). Carbon stocks in mixed planted forests were compared to carbon stocks in the average of constituent species monocultures, the best performing constituent monoculture (from a pool of 26 species) and, for a subset of comparisons, commercial species monocultures (from a pool of 11 species). Two species mixtures were the most well represented across the diversity range (species richness 2-6) (Fig. S2). Geographically the study sites used were biased to the northern hemisphere, with no sites in Africa or South America (Fig. S3). The age of study sites ranged from 3.5 to 28 years, with most comparisons from sites less than 20 years old (Fig. S1d).

Our overall meta-analyses showed that aboveground carbon stocks in mixed planted forests were higher than in the average of the monocultures (pooled SMD 1.41 [0.75, 2.07], k = 79, Fig. 1) and higher than in commercial monocultures (pooled SMD 1.06 [0.47, 1.65], k = 38, Fig. 1). Carbon stocks in mixed planted forests were on average higher than the most productive monoculture, but the confidence interval overlapped zero slightly (pooled SMD 0.62 [−0.0095, 1.25], k = 79, Fig. 1). We noted that for the comparison to the best monoculture, removal of the most extreme outliers (SMD >6 and <-3) altered the confidence interval for this result (pooled SMD 0.75 [0.32, 1.17], k = 73) (Supplement S2). Although the statistical tests are based on SMDs (in units of standard deviations), the underlying data suggest that mean carbon in the mixed treatments was 171% [95% CI 144, 198] (Table S3) of the carbon in the average of monocultures, 174% [95% CI 121, 127] of the carbon in commercial monocultures (Table S4) and 127% [95% CI 111, 142] of the carbon in the most productive monocultures (Table S3). The community with the greatest relative gain in aboveground carbon was a 3.5-year-old four-species mixture comprised of *Betula pendula, Fagus sylvatica, Quercus petraea* and *Tilia platyphyllos* in Saxony-Anhalt, Germany, which contained 43.9 Mg/ha of carbon compared to an average of 6.5 Mg/ha across the monocultures and 13.7 Mg/ha in the best monoculture (*Betula pendula*) (Table S3).

**Figure 1.**
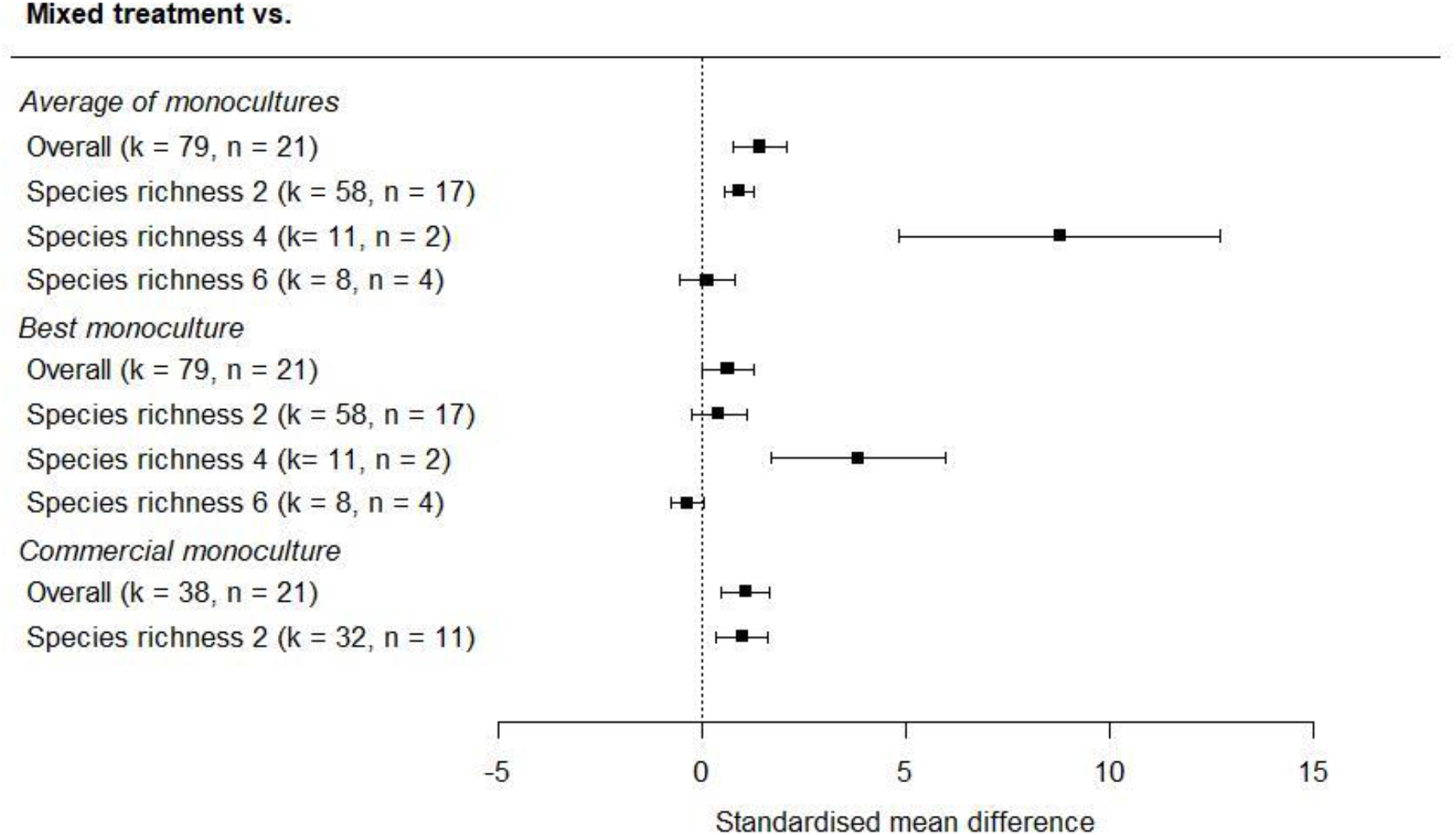
The effect of diversification of planted forests on aboveground carbon stocks relative to the average of associated monocultures, the best associated monoculture, and monocultures of commercial species, using all comparisons and subset by level of species richness. Effect sizes are standardised mean differences. Confidence intervals overlapping zero suggest no statistically detectible effect of diversification. Positive values indicate higher carbon stocks in mixtures than in monocultures. Number of comparisons (k) and number of study sites (n) are shown.

The greatest positive difference in carbon stocks was seen in mixed planted forests with four species, which had greater aboveground carbon stocks than the average of monocultures (pooled SMD 8.76 [95% CI 4.82, 12.7], k = 11, Fig. 1) and most productive monocultures (pooled SMD 3.82 [1.69, 5.95], k = 11, Fig. 1). Using the raw data, we calculated that carbon in the four-species mixtures was 411% [95% CI 325, 497] of the carbon in the average of monocultures and 232% [95%CI 218, 247] of the carbon in the best monoculture. Two-species mixtures also had greater aboveground carbon stocks than the average of monocultures (pooled SMD 0.90 [0.53, 1.27], k = 58, Fig. 1) and commercial monocultures (pooled SMD 0.97 [0.34, 1.61], k = 32, Fig. 1), but carbon stocks were not significantly greater than the most productive monocultures (pooled SMD 0.40 [−0.28, 1.08], k = 58, Fig. 1). Carbon in two-species mixtures was 135% [95% CI 122, 149] of carbon in the average of monocultures and 113% [95% CI 107, 120] of carbon in the most productive monoculture (Table S3). There was no clear difference in aboveground carbon stocks in six-species mixtures compared to the average of monocultures (pooled SMD 0.11 [−0.55, 0.77], k = 8, Fig. 1) and potential under-yielding relative to the most productive monoculture, although with a wide interval (pooled SMD –0.38 [–0.79, 0.042], k = 8, Fig. 1). When we fitted a moderator for species richness for the comparison to the average of monocultures and the best monoculture, we found that the moderator was significant for the comparison of mixtures to the average of monocultures (Q_M_ = 46.79, p < 0.01, k = 79, Fig. S4a), but not for the comparison to the best monoculture (Q_M_ 8.78, p = 0.067, k = 79, Fig. S4b).

Moderators for presence vs. absence of a nitrogen-fixer in mixtures (analyses possible for two-species mixtures only) were not significant for the comparison to the average of monocultures (Q_M_ = 0.37, p = 0.54, k = 58), best monoculture (Q_M_ = 1.35, p = 0.25, k = 58) or commercial species monoculture (Q_M_ = 1.33, p = 0.25, k = 32). We used data subset by presence or absence of a nitrogen-fixing species in the mixture to estimate the pooled effect sizes for aboveground carbon stocks in each group (Fig. 2a). In mixtures with a nitrogen-fixer present carbon was 145% [95% CI 136, 153] of the carbon in the average of monocultures and in mixtures without a nitrogen-fixer carbon was 128% [95% CI 120, 136] of the carbon in the average of monocultures.

**Figure 2.**
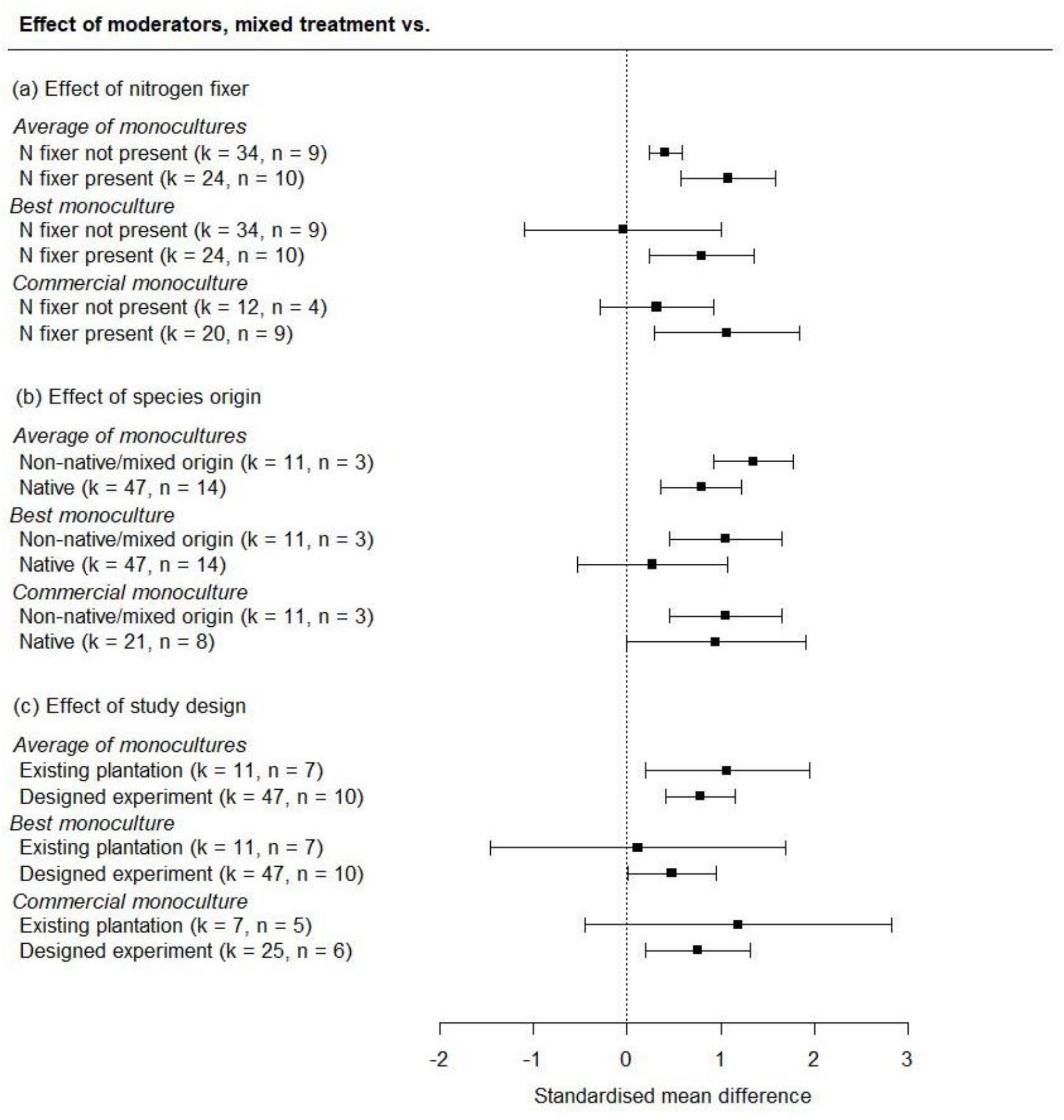
The effect of (a) diversification with/without a nitrogen-fixing species, (b) diversification with all native or non-native/mixed origin species, and (c) study design, on aboveground carbon stocks relative to the average of associated monocultures, the best associated monoculture, and monocultures of commercial species (analysis limited to two-species mixtures). Effect sizes are standardised mean differences, calculated using data subset by presence or absence of a nitrogenfixing species in the mixture, species origin, and study design. Confidence intervals overlapping zero suggest no statistically detectible effect of diversification. Positive values indicate higher carbon stocks in mixtures than in monocultures. Number of comparisons (k) and number of study sites (n) are shown.

Species origins in the mixed treatment (analyses possible for two-species mixtures only) were not significant for any of the comparisons (average of monocultures, Q_M_ = 0.94, p = 0.16, k = 58; best performing monoculture Q_M_ = 0.75, p = 0.39, k = 58; commercial monoculture Q_M_ = 0.091, p = 0.76, k = 32). Effect sizes estimated for each level of the moderator using subsets of the data confirmed that there was no clear effect of diversification based on the geographic origin of the species used (Fig. 2b). In mixtures of non-native species or a mixture of non-native and native species carbon was 156% [95% CI 148, 164] of the carbon in the average of monocultures and in mixtures with native species only carbon was 130% [95% CI 122, 137] of the carbon in the average of monocultures.

The effect of study type (analyses possible for two-species mixtures only) was not significant for any of the mixed to monoculture comparisons (average of monocultures Q_M_ = 0.24, p = 0.62, k = 58; best monoculture Q_M_ = 0.63, p = 0.43, k = 58; commercial species monoculture Q_M_ = 0.46, p = 0.50, k = 32). Subsequently, overall effect sizes were estimated using subsets of the data, confirming that there were no clear differences in the effect of diversification based on study type (Fig. 2c). In studies using forestry plantations, carbon in the mixed treatment was 128% [95% CI 120, 135] of the carbon in the average of monocultures and in experimental studies carbon in mixtures was 136% [95% CI 129, 144] of the carbon in the average of monocultures.

Analyses of the effect of age were possible for two-species mixtures only. Age had a better fit as a quadratic predictor rather than linear predictor for the comparison of mixed to the average of monocultures (AIC 155.9 vs. 162.6), showing a peak in the positive effect of diversification at the middle of the age range (quadratic term −0.013 [−0.023, −0.003]) (Fig. S5a). For the relationship between age and SMD relative to the best monoculture, a post-hoc quadratic predictor provided a better fit than a linear predictor (AIC 149.3 vs. 158.7), showing a peak in the positive effect of diversification before a downturn after age 17 yrs (quadratic term −0.019 [95% CI −0.029, −0.008]) (Fig. S5c). For the relationship between age and the difference in aboveground carbon stocks for mixed treatments compared to commercial monocultures a linear relationship provided the best fit but was not statistically distinguishable from zero (Q_M_ = 0.14, p = 0.71, k = 32, Fig. S5b).

## Discussion

Mixed planted forests clearly outperformed the average of monocultures and commercial monocultures, with no carbon stock penalty even relative to the best performing monoculture. This highlights the opportunity to maximise carbon stocks through diversification of the increasing area of planted forests worldwide (FAO, 2020). We found a peak in the diversity benefit in four-species mixtures. However, our meta-analysis also highlights the lack of data from planted forests at higher levels of tree diversity, and more data across the range of species richness is needed to better characterise this relationship. We noted that removal of the most extreme outliers altered the results for the comparison to the best monoculture, such that mixtures clearly outperformed the monoculture (Supplement S2). This further emphasises the need for more data to explore these trends. Given the young age of the planted forests used in our meta-analysis (3.5-28 years), and the expectation that diversity relationships will strengthen over time, further analysis as these forests age and from older planted forests would be informative (Guerrero-Ramírez et al., 2017). Alongside the expected benefits of increased provision of other ecosystem services, enhanced resilience, and resistance to pests and disease, our results further support diversifying planted forests (Aerts & Honnay, 2011; Messier et al., 2021). A critical next step is to integrate this information with analyses of the economics of diversifying plantations (Hildebrandt & Knoke, 2011).

The greatest positive effect of diversification was in mixtures with four species. Other studies have identified a hump-shaped (Xu et al., 2020), positive (Huang et al., 2018), and positive plateauing relationship (Liang et al., 2016; Zhang et al., 2012) between tree species richness and productivity in forest systems. However, in all these studies the relationship remained positive beyond a species richness of 20, except in Zhang et al. 2012, where the relationship plateaued at six species. Our data indicates an apparent peak in the positive effect of diversification in four species mixtures, however, all but one effect size for this species richness level is from one study, where the experimental plots were aged 3.5 yrs. Our dataset limited exploration of the diversity gradient, and the paucity of studies with higher levels of species richness could also explain the less clear effect of diversification in mixtures with >4 species.

The selection effect may explain the lack of transgressive overyielding found in our analysis, as when the monoculture species is particularly high yielding, it is harder for the mixture to outperform this. This can often be the case in commercial plantations, where species are usually selected for their fast growth rates, as well as attributes such as wood quality (C. L. C. Liu et al., 2018). Moreover, the benefits of diversification may become more apparent over the longer term and under future climate conditions, since mixed plantations may be more likely to maintain productivity even under perturbation (Osuri et al., 2020). As the study sites used in our analysis age, diversity effects may become more pronounced, and the relationship between diversity and function has been shown to become increasingly non-saturating over time (Guerrero-Ramírez et al., 2017; Reich et al., 2012).

When we included nitrogen-fixer as a moderator in our models, which was only possible for two-species mixtures, there was no clear effect. Presence or absence of species with particularly influential functional traits are expected to have a large effect on productivity (Tilman et al., 1997) and addition of a nitrogen-fixing tree species is a reasonably common method of forest diversification, aiming to reduce the need for fertiliser inputs (Marron & Epron, 2019; Richards et al., 2010; Temperton et al., 2007). However, other studies have also found that the presence of nitrogen-fixers in a mixture does not explain overyielding (Jactel et al., 2018). It has been shown that different species of nitrogen-fixers fix nitrogen at different rates and that their contribution of nitrogen fixation to a forest stand can change as forests age (Batterman et al., 2013, 2018). This may complicate the mechanism by which nitrogen-fixers contribute to overall forest productivity and accumulation of carbon stocks, which may not be captured in an analysis of presence vs. absence of a nitrogen-fixer in the mixture.

We found that non-native/mixed origin planted forests did not clearly outperform native planted forests. We expected that mixtures of native species only could lead to lower aboveground carbon stocks than mixtures including non-native species, which are often selected for their fast growth rates (Heryati et al., 2011). Fast-growing species such as *Eucalyptus* and *Pinus* make up a large proportion of planted forests, collectively comprising c.75% of the world’s commercial plantations, often in locations where these species are not native (Marron & Epron, 2019). However, in many of the studies used here, the fast growing species selected for plantations were native to the study location, for example *Eucalyptus globulus* in Australia (Bauhus et al., 2004).

We also found that the benefit of diversification held up in both highly controlled (e.g., experiments) and less controlled environments. It has often been questioned whether the influence of biodiversity on ecosystem function observed in controlled experiments is expressed in natural and managed ecosystems, where abiotic forcing and complex interactions occur (Duffy et al., 2017; Manning et al., 2019). Both operational forestry plantations and scientific tree diversity experiments are subject to some level of control relative to naturally established forests, with biodiversity experiments more often being more tightly controlled than forestry plantations. However, we found no clear influence of the study context on the effect of diversification on carbon stocks.

There are some limitations to our study, we have already highlighted the lack of representation across the diversity gradient and lack of higher diversity studies. The study sites contributing to our meta-analysis are also relatively young, aged between 3.5 and 28 years. We accounted for the differing ages of the planted forests in our analysis by including a random effect for study site; however, we also assessed the relationship between forest age and the carbon accumulation relative to monocultures for two-species mixtures. Previous meta-analyses have found that diversity effects increase over time (Cardinale et al., 2011), which has also been observed in forest systems (Guerrero-Ramírez et al., 2017; Huang et al., 2018). We found a peak in effect of diversity at 17 years old, however, our a-priori expectation was for a linear or saturating relationship (e.g. Reich et al., 2012; Thakur et al., 2015) and these quadratic models were fitted post-hoc following visual inspection of the data. The apparent downturn after age 17 is driven by the three comparisons at age 28, from two study sites. Given that the effects of complementarity between species are expected to strengthen over time (Fargione et al., 2007; Reich et al., 2012), the lack of data from older forests limits our potential to explore the consequences of diversification on timescales relevant to the forestry industry (Huang et al., 2018). Longer term data from the TreeDivNet experiments used here will be an important resource for further investigation of the effects of diversification on productivity in the future (Verheyen et al., 2016).

Data limitations also restricted our ability to assess the mechanisms behind differences in carbon accumulation, we were only able to assess moderators individually and for two-species mixtures only. We therefore cannot rule out collinearity between our moderators, although none of them were significant. Furthermore, there are other potential drivers of carbon accumulation that we did not explore, for example mixtures of tree species with complementary light interception strategies have been found to show overyielding (Williams et al., 2021). Establishment of future plantations using species that are selected because they are expected to be functionally complementary will be more informative than some earlier biodiversity experiments where species were selected entirely at random (Ebeling et al., 2014; X. Liu et al., 2018). Beyond species choices, other management decisions may influence carbon stocks, such as tree density. Moreover, there are different mechanisms that could be used to establish mixed plantations. On the larger scales at which plantations are typically established, diversification could be achieved through intermixing of species at the individual tree level or through compartments/zones comprising different species. The preferred option will depend on the scale at which diversity influences carbon accumulation, resilience and delivery of ecosystem services and trade-offs with practical constraints of planting, management and harvesting practices. If increased productivity depends on complementarity between individuals of different species then intermixing of species will be important, whereas resilience to extreme climatic events or delivery of certain services may occur at the landscape scale (Aquilué et al., 2020). These uncertainties call for studies exploring the potential scale-dependence of biodiversity effects on critical ecosystem services (Gonzalez et al., 2020) to inform the types of systems that can optimize carbon storage and other ecosystem services, while minimizing practical and financial constraints.

Our study focuses on aboveground carbon stocks only, which is typically the most easily measured forest carbon pool. However, belowground carbon stocks, in tree biomass and in the soil, can be important, with soil carbon stocks ranging from up to 90% of the total carbon stock in boreal forests to c.50% in tropical forests (Malhi et al., 2002). Previous studies have found a neutral or negative influence of diversity on belowground biomass. A recent study showed that, while mixtures showed overyielding aboveground, the belowground response was neutral, although the overall impact on carbon stocks remained significantly positive (Martin-Guay et al., 2020). If this result is replicated in other tree diversity studies, this emphasises that, by focussing on aboveground biomass, we may overestimate the impact of tree diversification on forest carbon accumulation (Martin-Guay et al., 2020). Further data on the impact of tree diversity on total ecosystem carbon stocks and an assessment of the responses of different carbon pools would therefore be valuable.

Diversification of plantations is one of the key actions recommended in Messier et al. 2021 and in a recent IPBES-IPCCC report on biodiversity and climate change, which aims to identify synergies and trade-offs between biodiversity protection and climate change mitigation and adaptation (Pörtner et al., 2021). The justification for use of mixed species plantations is that they will store more carbon, be more resilient to perturbation and provide greater support for associated biodiversity (Messier et al., 2021; Pörtner et al., 2021). Our study further supports the use of mixed species plantations as a method to increase carbon stocks and hence climate change mitigation (Beugnon et al., 2021). Accumulating evidence that diversification can increase carbon storage, resistance to perturbation (Hutchison et al., 2018), resilience to pests and disease (Jactel & Brockerhoff, 2007) and delivery of other ecosystem services (Gamfeldt et al., 2013), provides a strong justification for wider implementation of diversification in plantations (Messier et al., 2021). Even where mixtures do not provide substantial increases in carbon storage over monocultures (and we re-emphasise that we find no clear yield losses in mixtures relative to the best monoculture), they may nevertheless be desirable to increase levels of diversity of both the trees and associated organisms (Schuldt et al., 2018, 2019), as well as providing other potential benefits.

## Supporting information

Supplement S2 - publication bias

## Acknowledgements

E.W.’s PhD was funded by the Natural Environment Research Council NE/L002612/1, Oxford-NERC Doctoral Training Partnership in Environmental Research. The Bezos Earth Fund supported S.C.C-P.’s time on this work. N.E. and O.L. gratefully acknowledge the support by the German Centre for Integrative Biodiversity Research (iDiv) funded by the German Research Foundation (DFG–FZT 118, 202548816). The contribution of P.R. was supported by the U.S. NSF Biological Integration Institutes grant DBI-2021898.

## Author contributions

S.C.C-P. and A.H. conceived the study idea, and E.W. shaped the research focus. E.W. conducted the data extraction and analysis. J.K. gave advice on the analysis. N.E., O.F., D.G., J.S.H., H.J., C.M., C.M., C.M., A.P., W.C.P, C.P., and P.B.R. contributed data to the analysis. E.W. led the writing, alongside A.H and S.C.C-P. All authors commented on and approved the manuscript.

## References

Aalde, H., Gonzalez, P., Gytarsky, M., Krug, T., Kurz, W. A., Ogle, S., Raison, J., Schoene, D., & Ravindranath, N. (2006). Chapter 4 Forest Land. In IPCC Guidelines for National Greenhouse Gas Inventories. Volume 4: Agriculture, Forestry and Other Land Uses. Intergovernmental Panel on Climate Change.

Aerts, R., & Honnay, O. (2011). Forest restoration, biodiversity and ecosystem functioning. BMC Ecology, 11(29).

Ampoorter, E., Barbaro, L., Jactel, H., Baeten, L., Boberg, J., Carnol, M., Castagneyrol, B., Charbonnier, Y., Dawud, S. M., Deconchat, M., Smedt, P. De, Wandeler, H. De, Guyot, V., Hättenschwiler, S., Joly, F. X., Koricheva, J., Milligan, H., Muys, B., Nguyen, D.,… Allan, E. (2020). Tree diversity is key for promoting the diversity and abundance of forest-associated taxa in Europe. Oikos, 129(2), 133–146. https://doi.org/10.1111/oik.06290

Aquilué, N., Filotas, é., Craven, D., Fortin, M. J., Brotons, L., & Messier, C. (2020). Evaluating forest resilience to global threats using functional response traits and network properties. Ecological Applications, 30(5), 1–14. https://doi.org/10.1002/eap.2095

Batterman, S. A., Hall, J. S., Turner, B. L., Hedin, L. O., LaHaela Walter, J. K., Sheldon, P., & van Breugel, M. (2018). Phosphatase activity and nitrogen fixation reflect species differences, not nutrient trading or nutrient balance, across tropical rainforest trees. Ecology Letters, 21(10), 1486–1495. https://doi.org/10.1111/ele.13129

Batterman, S. A., Hedin, L. O., Van Breugel, M., Ransijn, J., Craven, D. J., & Hall, J. S. (2013). Key role of symbiotic dinitrogen fixation in tropical forest secondary succession. Nature, 502(7470), 224–227. https://doi.org/10.1038/nature12525

Bauhus, J., Van Winden, A. P., & Nicotra, A. B. (2004). Aboveground interactions and productivity in mixed-species plantations of Acacia mearnsii and Eucalyptus globulus. Canadian Journal of Forest Research, 34(3), 686–694. https://doi.org/10.1139/x03-243

Beugnon, R., Ladouceur, E., Sünnemann, M., Cesarz, S., & Eisenhauer, N. (2021). Diverse forests are cool: Promoting diverse forests to mitigate carbon emissions and climate change. Journal of Sustainable Agriculture and Environment. https://doi.org/10.1002/sae2.12005

Cardinale, B. J., Duffy, J. E., Gonzalez, A., Hooper, D. U., Perrings, C., Venail, P., Narwani, A., MacE, G. M., Tilman, D., Wardle, D. A., Kinzig, A. P., Daily, G. C., Loreau, M., Grace, J. B., Larigauderie, A., Srivastava, D. S., & Naeem, S. (2012). Biodiversity loss and its impact on humanity. Nature, 486(7401), 59–67. https://doi.org/10.1038/nature11148

Cardinale, B. J., Matulich, K. L., Hooper, D. U., Byrnes, J., Duffy, J. E., Gamfeldt, L., Balvanera, P., O’Connor, M. I., & Gonzalez, A. (2011). The functional role of producer diversity in ecosystems. American Journal of Botany, 98(3), 572–592.

Cook-Patton, S. C., & Agrawal, A. A. (2014). Exotic plants contribute positively to biodiversity functions but reduce native seed production and arthropod richness. Ecology, 95(6), 1642–1650. https://doi.org/10.1890/13-0782.1

Di Sacco, A., Hardwick, K., Blakesley, D., Brancalion, P. H. S., Breman, E., Rebola, L. C., Chomba, S., Dixon, K., Elliott, S., Ruyonga, G., Shaw, K., Smith, P., Smith, R. J., & Antonelli, A. (2021). Ten Golden Rules for Reforestation to Optimise Carbon Sequestration, Biodiversity Recovery and Livelihood Benefits. Global Change Biology, 27(7), 1328–1348. https://doi.org/10.1111/gcb.15498

Duffy, J. E., Godwin, C. M., & Cardinale, B. J. (2017). Biodiversity effects in the wild are common and as strong as key drivers of productivity. Nature, 549(7671), 261–264. https://doi.org/10.1038/nature23886

Ebeling, A., Pompe, S., Baade, J., Eisenhauer, N., Hillebrand, H., Proulx, R., Roscher, C., Schmid, B., Wirth, C., & Weisser, W. W. (2014). A trait-based experimental approach to understand the mechanisms underlying biodiversity-ecosystem functioning relationships. Basic and Applied Ecology, 15(3), 229–240. https://doi.org/10.1016/j.baae.2014.02.003

Ewel, J. J., Celis, G., & Schreeg, L. (2015). Steeply Increasing Growth Differential Between Mixture and Monocultures of Tropical Trees. Biotropica, 47(2), 162–171. https://doi.org/10.1111/btp.12190

FAO. (2020). Global Forest Resources Assessment 2020: Main report. https://doi.org/10.4324/9781315184487-1

Fargione, J., Tilman, D., Dybzinski, R., Lambers, J. H. R., Clark, C., Harpole, W. S., Knops, J. M. H., Reich, P. B., & Loreau, M. (2007). From selection to complementarity: Shifts in the causes of biodiversity-productivity relationships in a long-term biodiversity experiment. Proceedings of the Royal Society B: Biological Sciences, 274(1611), 871–876. https://doi.org/10.1098/rspb.2006.0351

Forster, E. J., Healey, J. R., Dymond, C., & Styles, D. (2021). Commercial afforestation can deliver effective climate change mitigation under multiple decarbonisation pathways. Nature Communications, 12, 3831. https://doi.org/10.1038/s41467-021-24084-x

Frey, C., Penman, J., Hanle, L., Monni, S., & Ogle, S. (2006). Chapter 3: Uncertainties. In 2006 IPCC Guidelines for National Greenhouse Gas Inventories. Volume 1: General Guidance and Reporting. https://doi.org/10.4324/9781315548906

Gamfeldt, L., Snäll, T., Bagchi, R., Jonsson, M., Gustafsson, L., Kjellander, P., Ruiz-Jaen, M. C., Fröberg, M., Stendahl, J., Philipson, C. D., Mikusiński, G., Andersson, E., Westerlund, B., Andrén, H., Moberg, F., Moen, J., & Bengtsson, J. (2013). Higher levels of multiple ecosystem services are found in forests with more tree species. Nature Communications, 4(1), 1340. https://doi.org/10.1038/ncomms2328

Gonzalez, A., Germain, R. M., Srivastava, D. S., Filotas, E., Dee, L. E., Gravel, D., Thompson, P. L., Isbell, F., Wang, S., Kéfi, S., Montoya, J., Zelnik, Y. R., & Loreau, M. (2020). Scaling-up biodiversity-ecosystem functioning research. Ecology Letters, 23(4), 757–776. https://doi.org/10.1111/ele.13456

Griscom, B. W., Adams, J., Ellis, P. W., Houghton, R. A., Lomax, G., Miteva, D. A., Schlesinger, W. H., Shoch, D., Siikamäki, J. V, Smith, P., Woodbury, P., Zganjar, C., Blackman, A., Campari, J., Conant, R. T., Delgado, C., Elias, P., Gopalakrishna, T., Hamsik, M. R.,… Fargione, J. (2017). Natural climate solutions. Proceedings of the National Academy of Sciences of the United States of America, 114(44), 11645–11650. https://doi.org/10.1073/pnas.1710465114

Guerrero-Ramírez, N. R., Craven, D., Reich, P. B., Ewel, J. J., Isbell, F., Koricheva, J., Parrotta, J. A., Auge, H., Erickson, H. E., Forrester, D. I., Hector, A., Joshi, J., Montagnini, F., Palmborg, C., Piotto, D., Potvin, C., Roscher, C., Van Ruijven, J., Tilman, D.,… Eisenhauer, N. (2017). Diversity-dependent temporal divergence of ecosystem functioning in experimental ecosystems. Nature Ecology and Evolution, 1(11), 1639–1642. https://doi.org/10.1038/s41559-017-0325-1

Hedges, L. V. (1981). Distribution Theory for Glass’s Estimator of Effect Size and Related Estimators. Journal of Educational Statistics, 6(2), 107–128.

Heryati, Y., Abdu, A., Mahat, M. N., Abdul-Hamid, H., Jusop, S., Majid, N. M., Heriansyah, I., & Ahmad, K. (2011). Assessing forest plantation productivity of exotic and indigenous species on degraded secondary forests. American Journal of Agricultural and Biological Science, 6(2), 201–208. https://doi.org/10.3844/ajabssp.2011.201.208

Hildebrandt, P., & Knoke, T. (2011). Investment decisions under uncertainty-A methodological review on forest science studies. Forest Policy and Economics, 13(1), 1–15. https://doi.org/10.1016/j.forpol.2010.09.001

Huang, Y., Chen, Y., Castro-Izaguirre, N., Baruffol, M., Brezzi, M., Lang, A., Li, Y., Härdtle, W., Von Oheimb, G., Yang, X., Liu, X., Pei, K., Both, S., Yang, B., Eichenberg, D., Assmann, T., Bauhus, J., Behrens, T., Buscot, F.,… Schmid, B. (2018). Impacts of species richness on productivity in a large-scale subtropical forest experiment. Science, 362(6410), 80–83. https://doi.org/10.1126/science.aat6405

Hutchison, C., Gravel, D., Guichard, F., & Potvin, C. (2018). Effect of diversity on growth, mortality, and loss of resilience to extreme climate events in a tropical planted forest experiment. Scientific Reports, 8(1), 1–10. https://doi.org/10.1038/s41598-018-33670-x

Jactel, H., Bauhus, J., Boberg, J., Bonal, D., Castagneyrol, B., Gardiner, B., Gonzalez-Olabarria, J. R., Koricheva, J., Meurisse, N., & Brockerhoff, E. G. (2017). Tree Diversity Drives Forest Stand Resistance to Natural Disturbances. Current Forestry Reports, 3(3), 223–243. https://doi.org/10.1007/s40725-017-0064-1

Jactel, H., & Brockerhoff, E. G. (2007). Tree diversity reduces herbivory by forest insects. Ecology Letters, 10(9), 835–848. https://doi.org/10.1111/j.1461-0248.2007.01073.x

Jactel, H., Gritti, E. S., Drössler, L., Forrester, D. I., Mason, W. L., Morin, X., Pretzsch, H., & Castagneyrol, B. (2018). Positive biodiversity–productivity relationships in forests: Climate matters. Biology Letters, 14(4), 12–15. https://doi.org/10.1098/rsbl.2017.0747

Jactel, H., Moreira, X., & Castagneyrol, B. (2021). Tree Diversity and Forest Resistance to Insect Pests: Patterns, Mechanisms, and Prospects. Annual Review of Entomology, 66, 277–296. https://doi.org/10.1146/annurev-ento-041720-075234

Jucker, T., Bouriaud, O., Avacaritei, D., & Coomes, D. A. (2014). Stabilizing effects of diversity on aboveground wood production in forest ecosystems: Linking patterns and processes. Ecology Letters, 17(12), 1560–1569. https://doi.org/10.1111/ele.12382

Jucker, T., Bouriaud, O., & Coomes, D. A. (2015). Crown plasticity enables trees to optimize canopy packing in mixed-species forests. Functional Ecology, 29(8), 1078–1086. https://doi.org/10.1111/1365-2435.12428

Lewis, S. L., Wheeler, C. E., Mitchard, E. T. A., & Koch, A. (2019). Restoring natural forests is the best way to remove atmospheric carbon. Nature, 568(7750), 25–28. https://doi.org/10.1038/d41586-019-01026-8

Liang, J., Crowther, T. W., Picard, N., Wiser, S., Zhou, M., Alberti, G., Schulze, E.-D., McGuire, A. D., Bozzato, F., Pretzsch, H., de-Miguel, S., Paquette, A., Hérault, B., Scherer-Lorenzen, M., Barrett, C. B., Glick, H. B., Hengeveld, G. M., Nabuurs, G.-J., Pfautsch, S.,… Reich, P. B. (2016). Positive biodiversity-productivity relationship predominant in global forests. Science, 354(6309), aaf8957. https://doi.org/10.1126/SCIENCE.AAF8957

Liu, C. L. C., Kuchma, O., & Krutovsky, K. V. (2018). Mixed-species versus monocultures in plantation forestry: Development, benefits, ecosystem services and perspectives for the future. In Global Ecology and Conservation (Vol. 15, p. e00419). Elsevier. https://doi.org/10.1016/j.gecco.2018.e00419

Liu, X., Trogisch, S., He, J.-S., Niklaus, P. A., Bruelheide, H., Tang, Z., Erfmeier, A., Scherer-Lorenzen, M., Pietsch, K. A., Yang, B., Kühn, P., Scholten, T., Huang, Y., Wang, C., Staab, M., Leppert, K. N., Wirth, C., Schmid, B., & Ma, K. (2018). Tree species richness increases ecosystem carbon storage in subtropical forests. Proceedings. Biological Sciences, 285(1885), 20181240. https://doi.org/10.1098/rspb.2018.1240

Loreau, M., & Hector, A. (2001). Partitioning selection and complementarity in biodiversity experiments. Nature, 412(6842), 72–76. http://www.biology.mcgill.ca/faculty/loreau/

Malhi, Y., Meir, P., & Brown, S. (2002). Forests, carbon and global climate. Philisophical Transactions of the Royal Society of London, 360, 1567–1591. https://doi.org/10.1098/rsta.2002.1020

Manning, P., Loos, J., Barnes, A. D., Batáry, P., Bianchi, F. J. J. A., Buchmann, N., De Deyn, G. B., Ebeling, A., Eisenhauer, N., Fischer, M., Fründ, J., Grass, I., Isselstein, J., Jochum, M., Klein, A. M., Klingenberg, E. O. F., Landis, D. A., Lepš, J., Lindborg, R.,… Tscharntke, T. (2019). Transferring biodiversity-ecosystem function research to the management of ‘real-world’ ecosystems. Advances in Ecological Research, 61, 323–356. https://doi.org/10.1016/BS.AECR.2019.06.009

Marron, N., & Epron, D. (2019). Are mixed-tree plantations including a nitrogen-fixing species more productive than monocultures? Forest Ecology and Management, 441(March), 242–252. https://doi.org/10.1016/j.foreco.2019.03.052

Martin-Guay, M. O., Paquette, A., Reich, P. B., & Messier, C. (2020). Implications of contrasted above- and below-ground biomass responses in a diversity experiment with trees. Journal of Ecology, 108(2), 405–414. https://doi.org/10.1111/1365-2745.13265

Mayoral, C., van Breugel, M., Cerezo, A., & Hall, J. S. (2017). Survival and growth of five Neotropical timber species in monocultures and mixtures. Forest Ecology and Management, 403, 1–11. https://doi.org/10.1016/J.FORECO.2017.08.002

McEwan, A., Marchi, E., Spinelli, R., & Brink, M. (2020). Past, present and future of industrial plantation forestry and implication on future timber harvesting technology. Journal of Forestry Research, 31(2), 339–351. https://doi.org/10.1007/s11676-019-01019-3

Messier, C., Bauhus, J., Sousa-Silva, R., Auge, H., Baeten, L., Barsoum, N., Bruelheide, H., Caldwell, B., Cavender-Bares, J., Dhiedt, E., Eisenhauer, N., Ganade, G., Gravel, D., Guillemot, J., Hall, J. S., Hector, A., Hérault, B., Jactel, H., Koricheva, J.,… Zemp, D. C. (2021). For the sake of resilience and multifunctionality, let’s diversify planted forests! Conservation Letters, July, 1–8. https://doi.org/10.1111/conl.12829

Nabuurs, G.-J., Verkerk, Pieter Johannes Schelhaas, M.-J., González Olabarria, José Ramón Trasobares, A., & Cienciala, E. (2018). Climate-Smart Forestry: mitigation impacts in three European regions. In Efi.Int (Issue April). https://www.efi.int/sites/default/files/files/publication-bank/2018/efi_fstp_6_2018.pdf

Osuri, A. M., Gopal, A., Raman, T. R. S., Defries, R., Cook-Patton, S. C., & Naeem, S. (2020). Greater stability of carbon capture in species-rich natural forests compared to species-poor plantations. Environmental Research Letters, 15(3). https://doi.org/10.1088/1748-9326/ab5f75

Pardos, M., del Río, M., Pretzsch, H., Jactel, H., Bielak, K., Bravo, F., Brazaitis, G., Defossez, E., Engel, M., Godvod, K., Jacobs, K., Jansone, L., Jansons, A., Morin, X., Nothdurft, A., Oreti, L., Ponette, Q., Pach, M., Riofrío, J.,… Calama, R. (2021). The greater resilience of mixed forests to drought mainly depends on their composition: Analysis along a climate gradient across Europe. Forest Ecology and Management, 481(October 2020), 118687. https://doi.org/10.1016/j.foreco.2020.118687

Piotto, D. (2008). A meta-analysis comparing tree growth in monocultures and mixed plantations. Forest Ecology and Management, 255(3–4), 781–786. https://doi.org/10.1016/J.FORECO.2007.09.065

Pörtner, H. O., Scholes, R. J., Agard, J., Archer, E., Arneth, A., Bai, X., Barnes, D., Burrows, M., Chan, L., Cheung, W. L., Diamond, S., Donatti, C., Duarte, C., Eisenhauer, N., Foden, W., Gasalla, M. A., Handa, C., Hickler, T., Hoegh-Guldberg, O.,… Ngo, H. T. (2021). IPBES-IPCC co-sponsored workshop report on biodiversity and climate change; IPBES and IPCC. https://doi.org/10.5281/zenodo.4782538

R Core Team. (2020). R: A Language and Environment for Statistical Computing. https://www.r-project.org/

Reich, P. B., Tilman, D., Isbell, F., Mueller, K., Hobbie, S. E., Flynn, D. F. B., & Eisenhauer, N. (2012). Impacts of biodiversity loss escalate through time as redundancy fades. Science, 336(6081), 589–592. https://doi.org/10.1126/science.1217909

Richards, A. E., Forrester, D. I., Bauhus, J., & Scherer-Lorenzen, M. (2010). The influence of mixed tree plantations on the nutrition of individual species: A review. Tree Physiology, 30(9), 1192–1208. https://doi.org/10.1093/treephys/tpq035

Schuldt, A., Assmann, T., Brezzi, M., Buscot, F., Eichenberg, D., Gutknecht, J., Härdtle, W., He, J. S., Klein, A. M., Kühn, P., Liu, X., Ma, K., Niklaus, P. A., Pietsch, K. A., Purahong, W., Scherer-Lorenzen, M., Schmid, B., Scholten, T., Staab, M.,… Bruelheide, H. (2018). Biodiversity across trophic levels drives multifunctionality in highly diverse forests. Nature Communications, 9(1). https://doi.org/10.1038/s41467-018-05421-z

Schuldt, A., Ebeling, A., Kunz, M., Staab, M., Guimarães-Steinicke, C., Bachmann, D., Buchmann, N., Durka, W., Fichtner, A., Fornoff, F., Härdtle, W., Hertzog, L. R., Klein, A. M., Roscher, C., Schaller, J., von Oheimb, G., Weigelt, A., Weisser, W., Wirth, C.,… Eisenhauer, N. (2019). Multiple plant diversity components drive consumer communities across ecosystems. Nature Communications, 10(1). https://doi.org/10.1038/s41467-019-09448-8

Seddon, N., Chausson, A., Berry, P., Girardin, C. A. J., Smith, A., & Turner, B. (2020). Understanding the value and limits of nature-based solutions to climate change and other global challenges. Philosophical Transactions of the Royal Society of London. Series B, Biological Sciences, 375(1794), 20190120. https://doi.org/10.1098/rstb.2019.0120

Temperton, V. M., Mwangi, P. N., Scherer-Lorenzen, M., Schmid, B., & Buchmann, N. (2007). Positive interactions between nitrogen-fixing legumes and four different neighbouring species in a biodiversity experiment. Oecologia, 151(2), 190–205. https://doi.org/10.1007/s00442-006-0576-z

Thakur, M. P., Milcu, A., Manning, P., Niklaus, P. A., Roscher, C., Power, S., Reich, P. B., Scheu, S., Tilman, D., Ai, F., Guo, H., Ji, R., Pierce, S., Ramirez, N. G., Richter, A. N., Steinauer, K., Strecker, T., Vogel, A., & Eisenhauer, N. (2015). Plant diversity drives soil microbial biomass carbon in grasslands irrespective of global environmental change factors. Global Change Biology, 21(11), 4076–4085. https://doi.org/10.1111/gcb.13011

Tilman, D., Isbell, F., & Cowles, J. M. (2014). Biodiversity and ecosystem functioning. Annual Review of Ecology, Evolution, and Systematics, 45, 471–493. https://doi.org/10.1146/annurev-ecolsys-120213-091917

Tilman, D., Knops, J., Wedin, D., Reich, P., Ritchie, M., & Siemann, E. (1997). The influence of functional diversity and composition on ecosystem processes. Science, 277(5330), 1300–1302. https://doi.org/10.1126/science.277.5330.1300

Tuck, S. L., O’Brien, M. J., Philipson, C. D., Saner, P., Tanadini, M., Dzulkifli, D., Godfray, H. C. J., Godoong, E., Nilus, R., Ong, R. C., Schmid, B., Sinun, W., Snaddon, J. L., Snoep, M., Tangki, H., Tay, J., Ulok, P., Wai, Y. S., Weilenmann, M.,… Hector, A. (2016). The value of biodiversity for the functioning of tropical forests: insurance effects during the first decade of the Sabah biodiversity experiment. Proceedings of the Royal Society B: Biological Sciences, 283(1844). https://doi.org/10.1098/rspb.2016.1451

van der Plas, F., Manning, P., Allan, E., Scherer-Lorenzen, M., Verheyen, K., Wirth, C., Zavala, M. A., Hector, A., Ampoorter, E., Baeten, L., Barbaro, L., Bauhus, J., Benavides, R., Benneter, A., Berthold, F., Bonal, D., Bouriaud, O., Bruelheide, H., Bussotti, F.,… Fischer, M. (2016). Jack-of-all-trades effects drive biodiversity–ecosystem multifunctionality relationships in European forests. Nature Communications, 7, 11109. https://doi.org/10.1038/ncomms11109

Verheyen, K., Vanhellemont, M., Auge, H., Baeten, L., Baraloto, C., Barsoum, N., Bilodeau-Gauthier, S., Bruelheide, H., Castagneyrol, B., Godbold, D., Haase, J., Hector, A., Jactel, H., Koricheva, J., Loreau, M., Mereu, S., Messier, C., Muys, B., Nolet, P.,… Scherer-Lorenzen, M. (2016). Contributions of a global network of tree diversity experiments to sustainable forest plantations. Ambio, 45(1), 29–41. https://doi.org/10.1007/s13280-015-0685-1

Viechtbauer, W. (2010). Conducting meta-analyses in {R} with the {metafor} package. Journal of Statistical Software, 36(3), 1–48. https://www.jstatsoft.org/v36/i03/

Williams, L. J., Butler, E. E., Cavender-Bares, J., Stefanski, A., Rice, K. E., Messier, C., Paquette, A., & Reich, P. B. (2021). Enhanced light interception and light use efficiency explain overyielding in young tree communities. Ecology Letters, 24(5), 996–1006. https://doi.org/10.1111/ele.13717

Williams, L. J., Paquette, A., Cavender-Bares, J., Messier, C., & Reich, P. B. (2017). Spatial complementarity in tree crowns explains overyielding in species mixtures. Nature Ecology and Evolution, 1(4). https://doi.org/10.1038/s41559-016-0063

Xu, S., Eisenhauer, N., Ferlian, O., Zhang, J., Zhou, G., Lu, X., Liu, C., & Zhang, D. (2020). Species richness promotes ecosystem carbon storage: Evidence from biodiversity-ecosystem functioning experiments. Proceedings of the Royal Society B: Biological Sciences, 287(1939). https://doi.org/10.1098/rspb.2020.2063rspb20202063

Zhang, Y., Chen, H. Y. H., & Reich, P. B. (2012). Forest productivity increases with evenness, species richness and trait variation: A global meta-analysis. Journal of Ecology, 100(3), 742–749. https://doi.org/10.1111/j.1365-2745.2011.01944.x

Zuppinger-Dingley, D., Schmid, B., Petermann, J. S., Yadav, V., De Deyn, G. B., & Flynn, D. F. B. (2014). Selection for niche differentiation in plant communities increases biodiversity effects. Nature, 515(7525), 108–111. https://doi.org/10.1038/nature13869

