## Supplement S1 - supplementary figures & tables for "Higher aboveground carbon stocks in mixed-species planted forests than monocultures – a meta-analysis"

**Supplement S1** – supplementary figures and tables

**Authors** Emily Warner[1], Susan C. Cook-Patton[2,3], Owen T. Lewis[4], Nick Brown[1], Julia Koricheva[5], Nico Eisenhauer[6,7], Olga Ferlian[6,7], Dominique Gravel[8], Jefferson S. Hall[9], Hervé Jactel[10], Carolina Mayoral[11,12], Céline Meredieu[10], Christian Messier[13,14], Alain Paquette[14], William C. Parker[15], Catherine Potvin[16,17], Peter B. Reich[18,19,20], Andy Hector[1]

**Affiliations**

[1]Department of Plant Sciences, University of Oxford, Oxford, UK

[2]The Nature Conservancy, Arlington, VA, USA

[3]Smithsonian Conservation Biology Institute, Front Royal, VA, USA

[4]Department of Zoology, University of Oxford, Oxford, UK

[5]Department of Biological Sciences, Royal Holloway University of London, UK

[6]German Centre for Integrative Biodiversity Research (iDiv) Halle-Jena-Leipzig, Leipzig, Germany

[7]Institute of Biology, Leipzig University, Leipzig, Germany

[8]Département de biologie, Université de Sherbrooke, Sherbooke, QC, Canada

[9]ForestGEO, Smithsonian Tropical Research Institute, Panama

[10]INRAE, Univ. Bordeaux, BIOGECO, F-33610 Cestas, France

[11] Birmingham Institute of Forest Research, Birmingham, UK

[12] School of Biosciences, Edgbaston Campus, University of Birmingham, UK

[13]Département de sciences naturelles and Institut des sciences de la forêt tempérée (ISFORT), Université du Québec en Outaouais (UQO), Ripon, QC, Canada

[14]Centre for Forest Research, Université du Québec à Montréal, PO Box 8888, Centre-ville Station, Montréal, QC H3C 3P8, Canada

[15]Ontario Forest Research Institute, Ontario Ministry of Northern Development, Mines, Natural Resources, and Forestry, Sault Ste. Marie, Ontario, Canada

[16]McGill University, Montréal, Québec, Canada

[17]Smithsonian Tropical Research Institute, Panama

[18]Department of Forest Resources, University of Minnesota, St. Paul, MN 55108, USA

[19] Hawkesbury Institute for the Environment, Western Sydney University, Penrith, NSW 2753, Australia

[20]Institute for Global Change Biology, and School for the Environment and Sustainability, University of Michigan, Ann Arbor, MI 48109, United States

**Contact information**

**Table S1** Information on the studies used in the meta-analysis, including whether they were obtained from the original literature search or from the Tree Diversity Network.

| | Study | Source | Site | Age | Country | Latitude | Longitude |
|---|---|---|---|---|---|---|---|
| **1** | (Debell et al., 1997) | Literature search | 1 | 11 | USA (Hawaii) | 19.860 | −155.150 |
| **2** | (Bauhus et al., 2004) | Literature search | 2 | 10 | Australia | −37.600 | 149.162 |
| **3** | (Forrester et al., 2013) | Literature search | 3 | 11 | Australia | −37.600 | 149.162 |
| **4** | (He et al., 2013) | Literature search | 5 | 28 | China | 22.0518 | 106.86854 |
| **5** | (Piotto et al., 2010) | Literature search | 6 | 17 | Costa Rica | 10.433333 | −83.983333 |
| **6** | (Piotto et al., 2010) | Literature search | 7 | 17 | Costa Rica | 10.433333 | −83.983333 |
| **7** | (Piotto et al., 2010) | Literature search | 8 | 16 | Costa Rica | 10.433333 | −83.983333 |
| **8** | (Nunes et al., 2014) | Literature search | 9 | 28 | Portugal | 41.400000 | −7.100000 |
| **9** | (Wang et al., 2009) | Literature search | 10 | 15 | China | 26.826522 | 109.859850 |
| **10** | (Mao et al., 2010) | Literature search | 11 | 5 | China | 41.705729 | 119.454350 |
| **11** | (Mao et al., 2010) | Literature search | 11 | 15 | China | 41.705729 | 119.454350 |
| **12** | (Nichols & Carpenter, 2006) | Literature search | 12 | 11 | Costa Rica | 9.000000 | −83.000000 |
| **13** | (Kaye et al., 2000) | Zhang *et al.* 2012 | 13 | 17 | USA (Hawaii) | 19.5 | -155.25 |
| **14** | (Son et al., 2007) | Zhang *et al.* 2012 | 14 | 27 | South Korea | 37.783333 | 127.8 |
| **15** | TreeDivNet – Orphee | TreeDivNet | 15 | 9 | France | 44.740100 | −0.796800 |
| **16** | TreeDivNet – Ident SSM | TreeDivNet | 16 | 6 | Canada | 46.546501 | −84.455650 |
| **17** | TreeDivNet – MyDiv | TreeDivNet | 17 | 3.5 | Germany | 51.383000 | 11.883000 |
| **18** | TreeDivNet – IDENT Auclair | TreeDivNet | 18 | 8 | Canada | 47.696611 | −68.656306 |
| **19** | TreeDivNet – IDENT Cloquet | TreeDivNet | 19 | 8 | Canada | 46.705083 | −92.524972 |

| | | | | | | | |
|---|---|---|---|---|---|---|---|
| **20** | TreeDivNet – Sardinilla | TreeDivNet | 20 | 16 | Panama | 9.316667 | −79.633333 |
| **21** | TreeDivNet – Agua Salud | TreeDivNet | 21 | 7 | Panama | 9.216667 | −79.7833333 |

**Table S2** Species classified as commercial species for the comparison of mixed plantation to monocultures of commercial species.

| Study | Country | Commercial species |
|---|---|---|
| Debell et al., 1997 | USA (Hawaii) | Eucalyptus saligna |
| Forrester et al., 2013 | Australia | Eucalyptus globulus |
| Piotto et al., 2010 | Costa Rica | Vochysia guatemalensis<br>Terminalia amazonia<br>Hyeronima alchorneoides |
| Nunes et al., 2014 | Portugal | Pseudotsuga menziesii |
| Wang et al., 2009 | China | Cunninghamia lanceolata |
| Mao et al., 2010 | China | Populus xiaozhuanica |
| Nichols & Carpenter, 2006 | Costa Rica | Terminalia amazonia |
| TreeDivNet – Agua Salud | Panama | Anacardium excelsum<br>Tabebuia rosea<br>Terminalia amazonia |
| Kaye et al., 2000 | USA (Hawaii) | Eucalyptus saligna |
| Son et al., 2007 | South Korea | Pinus koraiensis |

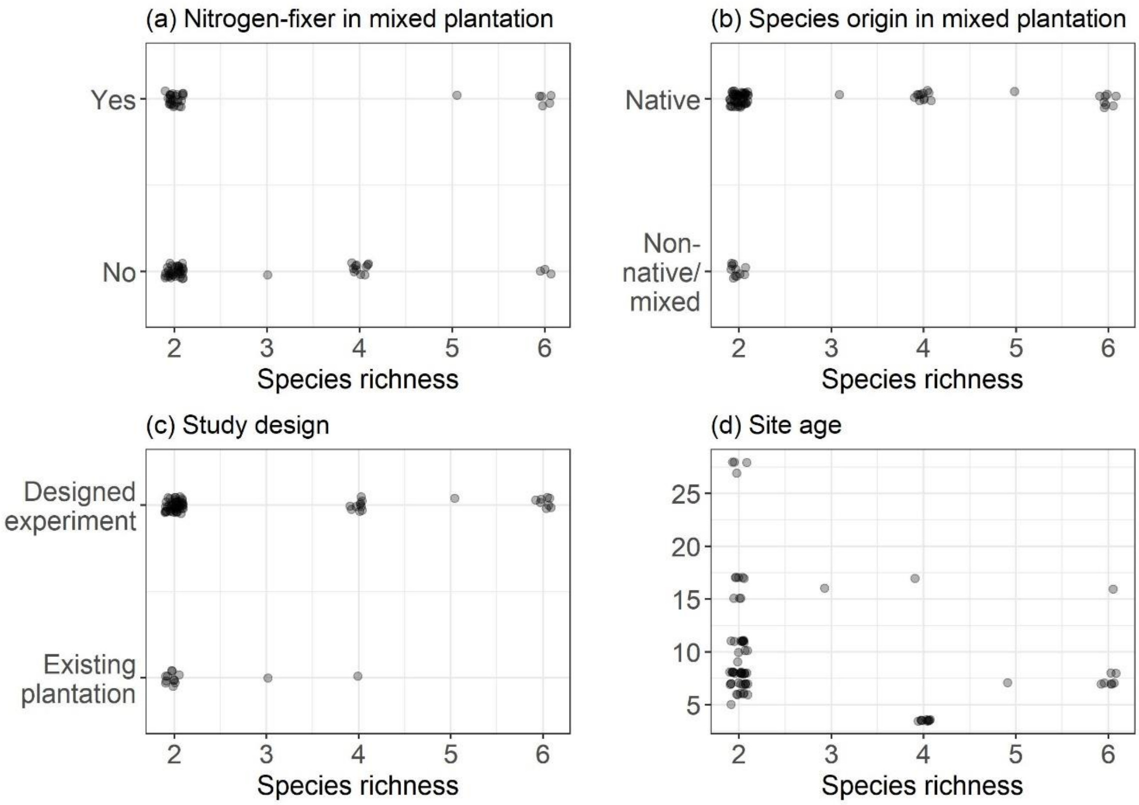


**Figure S1** Data representation for each moderator, (a) presence of nitrogen-fixer in the mixed plantation, (b) species origin of the species in the mixed plantation, (c) study design, (d) age of plantations, used to assess differences in carbon accumulation in mixed plantations compared to the average of monocultures/best monoculture.

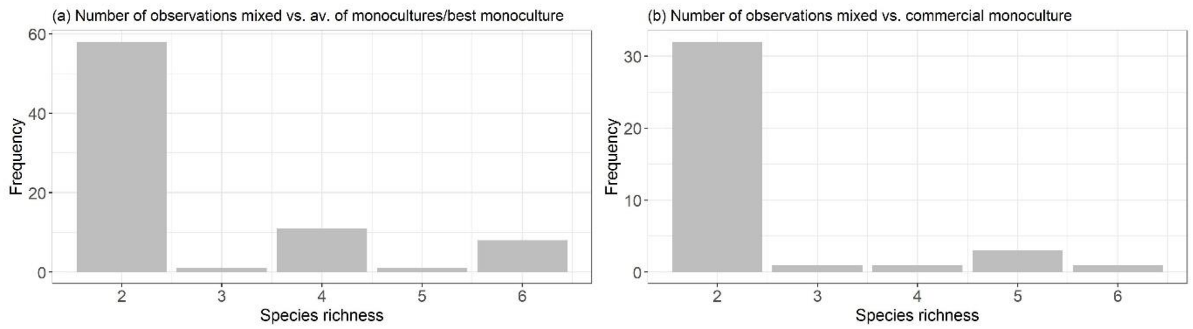


**Figure S2** Number of mixed to monoculture comparisons at each level of species richness for (a) mixed plantations compared to the average of relevant monocultures/best monoculture and (b) mixed plantations compared to commercial monocultures.

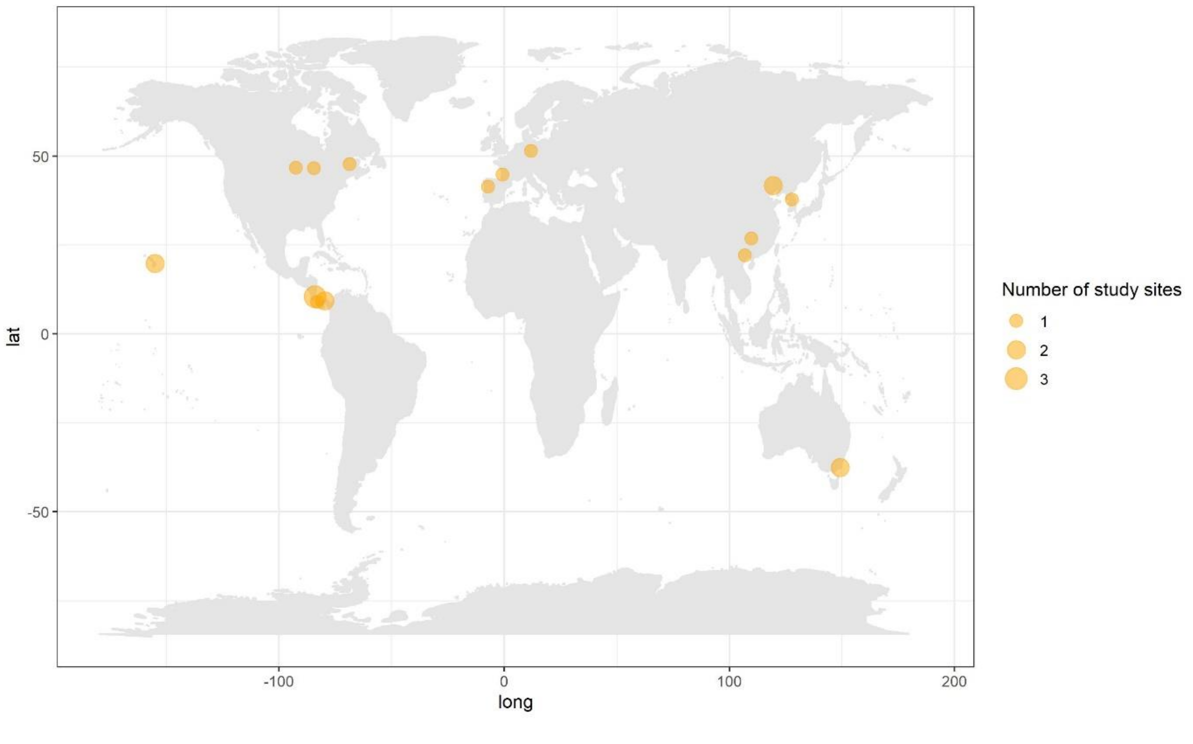


**Figure S3** The geographic distribution of study sites. Points are proportional to the number of independent sites in that location (where sites are spatially separate at a distance that would not be discernible at this map scale).

**Table S3** Carbon accumulation in each mixed plantation, the average of monoculture plantations and the most productive monoculture for each comparison. The % carbon in the mixed plantation relative to the average of monocultures and best monoculture is also shown. The comparisons are ordered by species richness and the % of the average of monocultures carbon in the mixed plantation. The age and species in the mixed and most productive monoculture plantations are given.

| SR | DP carbon (Mg/ha) | average of MPs carbon (Mg/ha) | most productive MP carbon (Mg/ha) | % of average of monocultures carbon in DP | % of most productive monoculture carbon in DP | Age | Mixed plantation species | Most productive monoculture species |
|---|---|---|---|---|---|---|---|---|
| 2 | 4.9 | 10.2 | 16.9 | 48 | 29 | 8 | *Larix laricina, Quercus rubra* | *Larix laricina* |
| 2 | 8.2 | 11.7 | 12.9 | 70 | 63 | 8 | *Acer saccharum, Picea glauca* | *Acer saccharum* |
| 2 | 9.9 | 13.6 | 16.9 | 73 | 58 | 8 | *Larix laricina, Pinus strobus* | *Larix laricina* |
| 2 | 68.5 | 82.2 | 82.2 | 83 | 83 | 27 | *Alnus hirsute, Pinus koraiensis* | *Pinus koraiensis* |
| 2 | 3.5 | 4.2 | 6.7 | 84 | 53 | 7 | *Anacardium excelsium, Tabebuia rosea* | *Anacardium excelsum* |
| 2 | 9.9 | 11.8 | 12.2 | 84 | 82 | 8 | *Betula papyrifera, Quercus rubra* | *Betula papyrifera* |
| 2 | 4.8 | 5.4 | 6.6 | 89 | 73 | 8 | *Acer saccharum, Picea glauca* | *Picea glauca* |
| 2 | 13.1 | 14.0 | 16.2 | 93 | 81 | 6 | *Pinus strobus, Quercus rubra* | *Pinus strobus* |
| 2 | 8.1 | 8.5 | 10.3 | 96 | 79 | 8 | *Picea glauca, Pinus strobus* | *Pinus strobus* |
| 2 | 7.0 | 6.9 | 7.0 | 102 | 100 | 7 | *Anacardium excelsium, Dalbergia retusa* | *Dalbergia retusa* |
| 2 | 9.2 | 9.0 | 9.0 | 102 | 102 | 5 | *Populus xiaozhuanica, Hippophae rhamnoides* | *Populus xiaozhuanica* |
| 2 | 67.2 | 63.8 | 88.3 | 105 | 76 | 28 | *Castanopsis hystrix, Pinus massoniana* | *Pinus massoniana* |
| 2 | 24.7 | 23.1 | 49.2 | 107 | 107 | 15 | *Cunninghamia lanceolata, Kalopanax septemlobus* | *Cunninghamia lanceolata* |
| 2 | 11.0 | 10.1 | 10.6 | 109 | 104 | 8 | *Picea glauca, Pinus strobus* | *Picea glauca* |
| 2 | 4.8 | 4.4 | 7.0 | 110 | 68 | 7 | *Dalbergia retusa, Tabebuia rosea* | *Dalbergia retusa* |
| 2 | 16.6 | 15.1 | 17.6 | 110 | 94 | 6 | *Larix larcinia, Picea glauca* | *Larix larcinia* |

| | | | | | | | | |
|---|---|---|---|---|---|---|---|---|
| 2 | 10.6 | 9.4 | 11.9 | 113 | 89 | 6 | *Acer saccharum, Quercus rubra* | *Quercus rubra* |
| 2 | 123.8 | 109.4 | 127.8 | 113 | 97 | 11 | *Eucalyptus saligna, Falcataria moluccana* | *Eucalyptus saligna* |
| 2 | 16.6 | 14.5 | 16.2 | 114 | 103 | 6 | *Betula papyrifera, Pinus strobus* | *Pinus strobus* |
| 2 | 12.3 | 10.6 | 10.8 | 117 | 114 | 8 | *Betula papyrifera, Pinus strobus* | *Betula papyrifera* |
| 2 | 5.2 | 4.4 | 6.7 | 118 | 78 | 7 | *Anacardium excelsium, Pachira quinata* | *Anacardium excelsum* |
| 2 | 12.9 | 10.8 | 12.2 | 119 | 106 | 8 | *Betula papyrifera, Pinus strobus* | *Betula papyrifera* |
| 2 | 43.0 | 36.0 | 36.0 | 119 | 119 | 11 | *Eucalyptus globulus, Acacia mearnsii* | *Acacia mearnsii* |
| 2 | 43.1 | 36.0 | 36.0 | 120 | 120 | 11 | *Eucalyptus globulus, Acacia mearnsii* | *Acacia mearnsii* |
| 2 | 131.9 | 109.4 | 127.8 | 120 | 103 | 11 | *Eucalyptus saligna, Falcataria moluccana* | *Eucalyptus saligna* |
| 2 | 60.0 | 49.2 | 49.2 | 122 | 122 | 15 | *Cunninghamia lanceolata, Alnus cremastogyne* | *Cunninghamia lanceolata* |
| 2 | 14.8 | 12.1 | 14.8 | 122 | 101 | 8 | *Larix laricina, Pinus strobus* | *Larix laricina* |
| 2 | 12.2 | 9.9 | 12.9 | 124 | 95 | 6 | *Acer saccharum, Betula papyrifera* | *Betula papyrifera* |
| 2 | 16.9 | 13.6 | 16.9 | 124 | 100 | 8 | *Larix laricina, Pinus strobus* | *Larix laricina* |
| 2 | 18.0 | 14.3 | 16.2 | 125 | 111 | 6 | *Picea glauca, Pinus strobus* | *Pinus strobus* |
| 2 | 10.8 | 8.5 | 10.3 | 128 | 105 | 8 | *Picea glauca, Pinus strobus* | *Pinus strobus* |
| 2 | 15.5 | 11.9 | 22.2 | 130 | 70 | 7 | *Terminalia amazonia, Tabebuia rosea* | *Terminalia amazonia* |
| 2 | 47.0 | 36.0 | 36.0 | 130 | 130 | 11 | *Eucalyptus globulus, Acacia mearnsii* | *Acacia mearnsii* |
| 2 | 19.1 | 14.6 | 22.2 | 131 | 86 | 7 | *Dalbergia retusa, Terminalia amazonia* | *Terminalia amazonia* |
| 2 | 144.8 | 109.4 | 127.8 | 132 | 113 | 11 | *Eucalyptus saligna, Falcataria moluccana* | *Eucalyptus saligna* |
| 2 | 17.4 | 13.1 | 14.8 | 133 | 118 | 8 | *Larix laricina, Quercus rubra* | *Larix laricina* |
| 2 | 31.4 | 23.5 | 30.6 | 134 | 103 | 10 | *Eucalyptus globulus, Acacia mearnsii* | *Acacia mearnsii* |
| 2 | 66.9 | 48.9 | 62.7 | 137 | 107 | 28 | *Pseudotsuga menziesii, Castanea sativa* | *Pseudotsuga menziesii* |

| | | | | | | | | |
|---|---|---|---|---|---|---|---|---|
| 2 | 19.9 | 14.4 | 22.2 | 138 | 90 | 7 | *Anacardium excelsium, Terminialia amazonia* | *Terminalia amazonia* |
| 2 | 17.4 | 12.5 | 12.9 | 139 | 135 | 8 | *Acer saccharum, Betula papyrifera* | *Acer saccharum* |
| 2 | 170.3 | 120.5 | 120.5 | 141 | 141 | 15 | *Populus xiaozhuanica, Hippophae rhamnoides* | *Populus xiaozhuanica* |
| 2 | 162.1 | 109.4 | 127.8 | 148 | 127 | 11 | *Eucalyptus saligna, Falcataria moluccana* | *Eucalyptus saligna* |
| 2 | 35.0 | 23.5 | 30.6 | 149 | 115 | 10 | *Eucalyptus globulus, Acacia mearnsii* | *Acacia mearnsii* |
| 2 | 35.4 | 23.5 | 30.6 | 151 | 116 | 10 | *Eucalyptus globulus, Acacia mearnsii* | *Acacia mearnsii* |
| 2 | 156.1 | 101.1 | 100.7 | 154 | 155 | 17 | *Eucalyptus saligna, Falcataria moluccana* | *Eucalyptus saligna* |
| 2 | 19.2 | 12.1 | 22.2 | 158 | 86 | 7 | *Pachira quinate, Terminalia amazonia* | *Terminalia amazonia* |
| 2 | 180.6 | 109.4 | 127.8 | 165 | 141 | 11 | *Eucalyptus saligna, Falcataria moluccana* | *Eucalyptus saligna* |
| 2 | 46.7 | 27.5 | 40.6 | 170 | 115 | 17 | *Vochysia guatemalensis, Jacaranda copaia* | *Vochysia guatemalensis* |
| 2 | 83.7 | 48.9 | 62.7 | 171 | 133 | 28 | *Pseudotsuga menziesii, Castanea sativa* | *Pseudotsuga menziesii* |
| 2 | 12.9 | 7.5 | 10.8 | 172 | 119 | 8 | *Acer saccharum, Betula papyrifera* | *Betula papyrifera* |
| 2 | 177.5 | 101.1 | 100.7 | 176 | 176 | 17 | *Eucalyptus saligna, Falcataria moluccana* | *Eucalyptus saligna* |
| 2 | 8.1 | 4.6 | 7.0 | 177 | 115 | 7 | *Dalbergia retusa, Pachira quinata* | *Dalbergia retusa* |
| 2 | 3.5 | 1.9 | 2.1 | 185 | 166 | 7 | *Pachira quinate, Tabebuia rosea* | *Pachira quinata* |
| 2 | 191.1 | 101.1 | 100.7 | 189 | 190 | 17 | *Eucalyptus saligna, Falcataria moluccana* | *Eucalyptus saligna* |
| 2 | 209.9 | 101.1 | 100.7 | 208 | 208 | 17 | *Eucalyptus saligna, Falcataria moluccana* | *Eucalyptus saligna* |
| 2 | 17.9 | 7.2 | 10.8 | 250 | 165 | 8 | *Betula papyrifera, Quercus rubra* | *Betula papyrifera* |
| 2 | 31.2 | 11.0 | 12.5 | 285 | 250 | 9 | *Betula pendula, Pinus pinaster* | *Pinus pinaster* |

| | | | | | | | | |
|---|---|---|---|---|---|---|---|---|
| 2 | 10.8 | 3.7 | 3.7 | 293 | 293 | 11 | *Terminalia amazonia, Inga edulis* | *Terminalia amazonia* |
| 3 | 26.5 | 18.5 | 22.1 | 144 | 120 | 16 | *Hyeronima alchorneoides, Vochysia ferruginea, Balizia elegans* | *Hyeronima alchorneoides* |
| 4 | 63.4 | 42.8 | 55.0 | 148 | 115 | 17 | *Terminalia amazonia, Virola koschnyi, Dipteryx panamensis* | *Terminalia amazonia* |
| 4 | 31.1 | 10.1 | 16.5 | 308 | 189 | 3.5 | *Aesculus hippocastanum, Fraxinus excelsior, Prunus avium, Sorbus aucuparia* | *Sorbus aucuparia* |
| 4 | 44.9 | 13.3 | 30.5 | 336 | 147 | 3.5 | *Betula pendula, Carpinus betulus, Quercus petraea, Tilia platyphyllos* | *Carpinus betulus* |
| 4 | 47.2 | 13.4 | 30.5 | 351 | 155 | 3.5 | *Betula pendula, Carpinus betulus, Fagus sylvatica, Tilia platyphyllos* | *Carpinus betulus* |
| 4 | 45.6 | 12.5 | 30.5 | 365 | 150 | 3.5 | *Betula pendula, Carpinus betulus, Fagus sylvatica, Quercus petraea* | *Carpinus betulus* |
| 4 | 51.6 | 14.1 | 16.9 | 367 | 304 | 3.5 | *Acer pseudoplatanus, Fraxinus excelsior, Prunus avium, Sorbus aucuparia* | *Acer pseudoplatanus* |
| 4 | 55.7 | 12.8 | 16.9 | 435 | 329 | 3.5 | *Acer pseudoplatanus, Aesculus hippocastanum, Fraxinus excelsior, Sorbus aucuparia* | *Acer pseudoplatanus* |
| 4 | 46.1 | 10.2 | 16.9 | 450 | 272 | 3.5 | *Acer pseudoplatanus, Aesculus hippocastanum, Fraxinus excelsior, Prunus avium* | *Acer pseudoplatanus* |
| 4 | 64.0 | 11.8 | 16.9 | 543 | 378 | 3.5 | *Acer pseudoplatanus, Aesculus hippocastanum, Prunus avium, Sorbus aucuparia* | *Acer pseudoplatanus* |
| 4 | 58.7 | 10.7 | 30.5 | 548 | 193 | 3.5 | *Carpinus betulus, Fagus sylvatica, Quercus petraea, Tilia platyphyllos* | *Carpinus betulus* |
| 4 | 43.9 | 6.5 | 13.7 | 674 | 321 | 3.5 | *Betula pendula, Fagus sylvatica, Quercus petraea, Tilia platyphyllos* | *Betula pendula* |

| | | | | | | | | |
|---|---|---|---|---|---|---|---|---|
| 5 | 14.4 | 7.9 | 22.2 | 182 | 65 | 7 | *Terminalia amazonia, Tabebuia rosea, Dalbergia retusa, Pachira quinate, Anacardium excelsium* | *Terminalia amazonia* |
| 6 | 2.6 | 6.7 | 6.7 | 39 | 39 | 7 | *Anacardium excelsium, Erythrina fusca, Gliricidia sepium, Inga punctata, Luehea speciose, Ochroma pyramidale* | *Anacardium excelsum* |
| 6 | 5.1 | 7.0 | 7.0 | 72 | 72 | 7 | *Dalbergia retusa, Erythrina fusca, Gliricidia sepium, Inga punctata, Luehea speciose, Ochroma pyramidale* | *Dalbergia retusa* |
| 6 | 17.2 | 22.2 | 22.2 | 78 | 78 | 7 | *Terminalia amazonia, Erythrina fusca, Gliricidia sepium, Inga punctata, Luehea speciose, Ochroma pyramidale* | *Terminalia amazonia* |
| 6 | 7.9 | 8.7 | 16.9 | 91 | 47 | 8 | *Acer saccharum, Betula papyrifera, Larix laricina, Picea glauca, Pinus strobus, Quercus rubra* | *Larix larici* |
| 6 | 1.7 | 1.7 | 1.7 | 102 | 102 | 7 | *Tabebuia rosea, Erythrina fusca, Gliricidia sepium, Inga punctata, Luehea speciose, Ochroma pyramidale* | *Tabebuia rosea* |
| 6 | 13.5 | 11.9 | 14.8 | 113 | 91 | 8 | *Acer saccharum, Betula papyrifera, Larix laricina, Picea glauca, Pinus strobus, Quercus rubra* | *Larix larici* |
| 6 | 36.9 | 23.3 | 36.3 | 158 | 102 | 16 | *Anacardium excelsum, Cedrela odorata, Hura crepitans, Luehea seemanii, Tabebuia rosea, Cordia alliodora* | *Luehea seemanii* |

| | | | | | | | | |
|---|---|---|---|---|---|---|---|---|
| 6 | 5.6 | 2.1 | 2.1 | 263 | 263 | 7 | *Pachira quinate, Erythrina fusca, Gliricidia sepium, Inga punctata, Luehea speciose, Ochroma pyramidale* | *Pachira quinata* |
| **Overall** | | | | **170.3 [95% CI 143.9, 196.8]** | **128.0 [95% CI 112.4, 143.5]** | | | |

**Table S4** Carbon accumulation in mixed plantations compared to commercial species monocultures. The % carbon in the mixed plantation relative to the commercial species monoculture is shown. The comparisons are ordered by species richness and the % of the commercial species monoculture carbon in the mixed plantation. The age and species in the mixed and commercial species monoculture plantations are given.

| SR | DP carbon (Mg/ha) | commercial MP carbon (Mg/ha) | % of commercial monoculture carbon in DP | Age | Mixed plantation species | Commercial monoculture species |
|---|---|---|---|---|---|---|
| 2 | 3.5 | 6.7 | 53 | 7 | Anacardium excelsium Tabebuia rosea | Anacardium excelsum |
| 2 | 15.5 | 22.2 | 70 | 7 | Terminalia amazonia Tabebuia rosea | Terminalia amazonia |
| 2 | 5.2 | 6.7 | 78 | 7 | Anacardium excelsium Pachira quinata | Anacardium excelsum |
| 2 | 68.5 | 82.2 | 83 | 27 | Alnus hirsuta Pinus koraiensis | Pinus koraiensis |
| 2 | 19.1 | 22.2 | 86 | 7 | Dalbergia retusa Terminalia amazonia | Terminalia amazonia |
| 2 | 19.2 | 22.2 | 86 | 7 | Pachira quinata Terminalia amazonia | Terminalia amazonia |
| 2 | 19.9 | 22.2 | 90 | 7 | Anacardium excelsium Terminialia amazonia | Terminalia amazonia |
| 2 | 123.8 | 127.8 | 97 | 11 | Eucalyptus saligna Falcataria moluccana | Eucalyptus saligna |
| 2 | 9.2 | 9.0 | 102 | 5 | Populus xiaozhuanica Hippophae rhamnoides | Populus xiaozhuanica |
| 2 | 131.9 | 127.8 | 103 | 11 | Eucalyptus saligna Falcataria moluccana | Eucalyptus saligna |
| 2 | 7.0 | 6.7 | 105 | 7 | Anacardium excelsium Dalbergia retusa | Anacardium excelsum |
| 2 | 66.9 | 62.7 | 107 | 28 | Pseudotsuga menziesii Castanea sativa | Pseudotsuga menziesii |
| 2 | 52.7 | 49.2 | 107 | 15 | Cunninghamia lanceolata Kalopanax septemlobus | Cunninghamia lanceolata |
| 2 | 144.8 | 127.8 | 113 | 11 | Eucalyptus saligna Falcataria moluccana | Eucalyptus saligna |
| 2 | 46.7 | 40.6 | 115 | 17 | Vochysia guatemalensis Jacaranda copaia | Vochysia guatemalensis |
| 2 | 60.0 | 49.2 | 122 | 15 | Cunninghamia lanceolata Alnus cremastogyne | Cunninghamia lanceolata |

| | | | | | | |
|---|---|---|---|---|---|---|
| 2 | 162.1 | 127.8 | 127 | 11 | Eucalyptus saligna Falcataria moluccana | Eucalyptus saligna |
| 2 | 83.7 | 62.7 | 133 | 28 | Pseudotsuga menziesii Castanea sativa | Pseudotsuga menziesii |
| 2 | 180.6 | 127.8 | 141 | 11 | Eucalyptus saligna Falcataria moluccana | Eucalyptus saligna |
| 2 | 170.3 | 120.5 | 141 | 15 | Populus xiaozhuanica Hippophae rhamnoides | Populus xiaozhuanica |
| 2 | 156.1 | 100.7 | 155 | 17 | Eucalyptus saligna Falcataria moluccana | Eucalyptus saligna |
| 2 | 177.5 | 100.7 | 176 | 17 | Eucalyptus saligna Falcataria moluccana | Eucalyptus saligna |
| 2 | 191.1 | 100.7 | 190 | 17 | Eucalyptus saligna Falcataria moluccana | Eucalyptus saligna |
| 2 | 31.4 | 16.5 | 191 | 10 | Eucalyptus globulus Acacia mearnsii | Eucalyptus globulus |
| 2 | 209.9 | 100.7 | 208 | 17 | Eucalyptus saligna Falcataria moluccana | Eucalyptus saligna |
| 2 | 3.5 | 1.7 | 209 | 7 | Pachira quinata Tabebuia rosea | Tabebuia rosea |
| 2 | 35.0 | 16.5 | 213 | 10 | Eucalyptus globulus Acacia mearnsii | Eucalyptus globulus |
| 2 | 35.4 | 16.5 | 215 | 10 | Eucalyptus globulus Acacia mearnsii | Eucalyptus globulus |
| 2 | 4.8 | 1.7 | 282 | 7 | Dalbergia retusa Tabebuia rosea | Tabebuia rosea |
| 2 | 10.8 | 3.7 | 293 | 11 | Terminalia amazonia Inga edulis | Terminalia amazonia |
| 2 | 19.9 | 6.7 | 298 | 7 | Anacardium excelsium Terminialia amazonia | Anacardium excelsum |
| 2 | 15.5 | 1.7 | 913 | 7 | Terminalia amazonia Tabebuia rosea | Tabebuia rosea |
| 3 | 26.5 | 22.1 | 120 | 16 | Hyeronima alchorneoides Vochysia ferruginea Balizia elegans | Hyeronima alchorneoides |
| 4 | 63.4 | 55.0 | 115 | 17 | Terminalia amazonia Virola koschnyi Dipteryx panamensis | Terminalia amazonia |
| 5 | 14.4 | 6.7 | 216 | 7 | Terminalia amazonia Tabebuia rosea Dalbergia retusa Pachira quinata Anacardium excelsium | Anacardium excelsum |
| 5 | 14.4 | 22.2 | 65 | 7 | Terminalia amazonia Tabebuia rosea Dalbergia retusa Pachira quinata Anacardium excelsium | Terminalia amazonia |

| | | | | | | |
|---|---|---|---|---|---|---|
| 5 | 14.4 | 1.7 | 851 | 7 | Terminalia amazonia Tabebuia rosea Dalbergia retusa Pachira quinata Anacardium excelsium | Tabebuia rosea |
| 6 | 17.2 | 22.2 | 78 | 7 | Terminalia amazonia Erythrina fusca Gliricidia sepium Inga punctata Luehea speciose Ochroma pyramidale | Terminalia amazonia |
| **Overall** | | | **176.6 [95% CI 119.4, 233.8]** | | | |

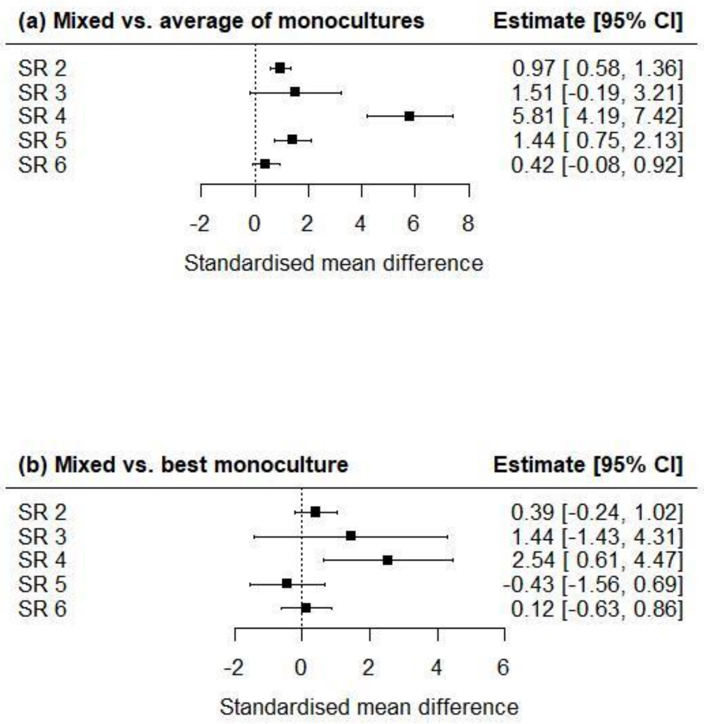


**Figure S4** The effect of diversification at different levels of species richness on aboveground carbon stocks relative to (a) the average of associated monocultures and (b) the best associated monocultures. Effect sizes are standardised mean differences, predicted from models with species richness fitted as a discrete moderator. Confidence intervals overlapping zero suggest no statistically detectible effect of diversification. Positive values indicate higher carbon stocks in mixtures than in monocultures.

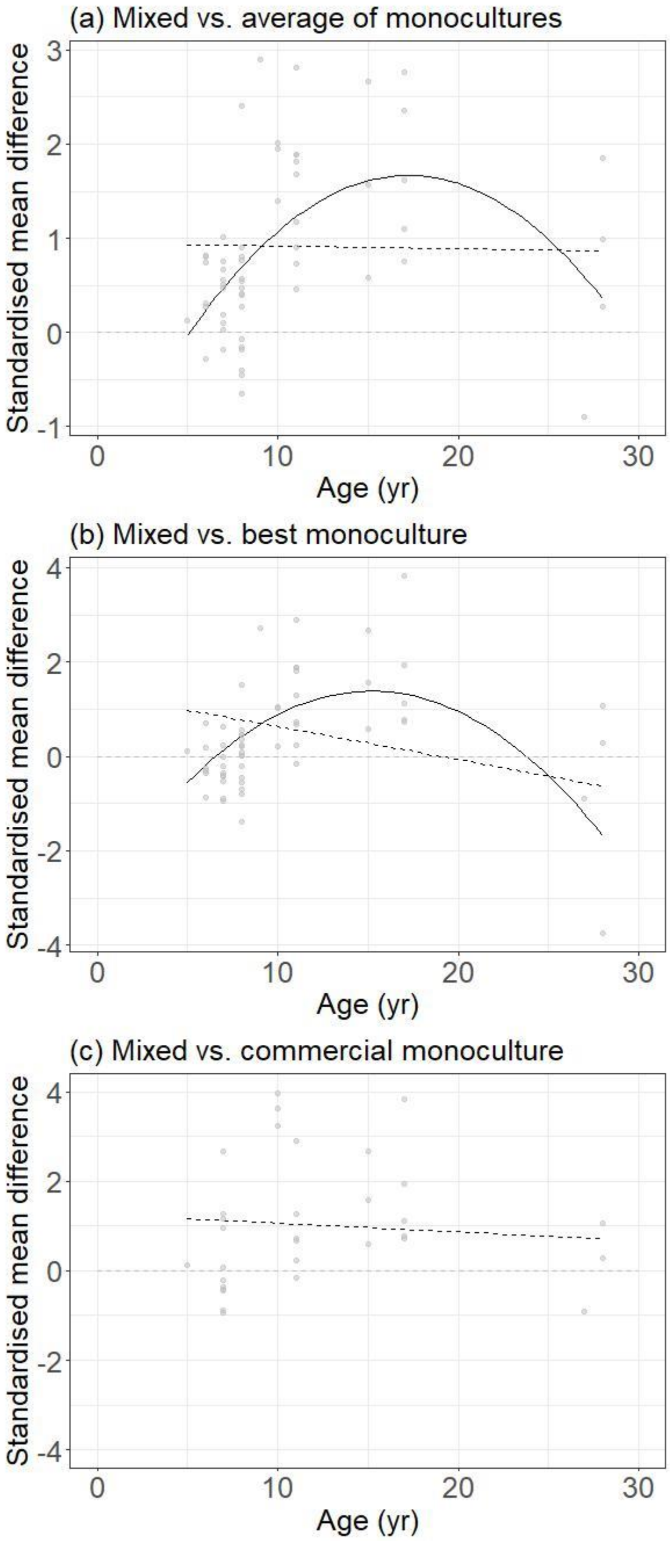


**Figure S5** The relationship between stand age and the standardised mean difference in aboveground carbon stocks (analysis limited to two-species mixtures) relative to (a) the average of relevant monocultures, k = 58; (b) the best associated monoculture, k = 58; (c) commercial monocultures, k = 32. Non-significant trends shown as dashed lines.
