## Supplement S2 - publication bias for "Higher aboveground carbon stocks in mixed-species planted forests than monocultures – a meta-analysis"

**Supplement S2** – Publication bias and removal of outliers

**Authors** Emily Warner[1], Susan C. Cook-Patton[2,3], Owen T. Lewis[4], Nick Brown[1], Julia Koricheva[5], Nico Eisenhauer[6,7], Olga Ferlian[6,7], Dominique Gravel[8], Jefferson S. Hall[9], Hervé Jactel[10], Carolina Mayoral[11,12], Céline Meredieu[10], Christian Messier[13,14], Alain Paquette[14], William C. Parker[15], Catherine Potvin[16,17], Peter B. Reich[18,19,20], Andy Hector[1]

**Affiliations**

[1]Department of Plant Sciences, University of Oxford, Oxford, UK

[2]The Nature Conservancy, Arlington, VA, USA

[3]Smithsonian Conservation Biology Institute, Front Royal, VA, USA

[4]Department of Zoology, University of Oxford, Oxford, UK

[5]Department of Biological Sciences, Royal Holloway University of London, UK

[6]German Centre for Integrative Biodiversity Research (iDiv) Halle-Jena-Leipzig, Leipzig, Germany

[7]Institute of Biology, Leipzig University, Leipzig, Germany

[8]Département de biologie, Université de Sherbrooke, Sherbooke, QC, Canada

[9]ForestGEO, Smithsonian Tropical Research Institute, Panama

[10]INRAE, Univ. Bordeaux, BIOGECO, F-33610 Cestas, France

[11] Birmingham Institute of Forest Research, Birmingham, UK

[12] School of Biosciences, Edgbaston Campus, University of Birmingham, UK

[13]Département de sciences naturelles and Institut des sciences de la forêt tempérée (ISFORT), Université du Québec en Outaouais (UQO), Ripon, QC, Canada

[14]Centre for Forest Research, Université du Québec à Montréal, PO Box 8888, Centre-ville Station, Montréal, QC H3C 3P8, Canada

[15]Ontario Forest Research Institute, Ontario Ministry of Northern Development, Mines, Natural Resources, and Forestry, Sault Ste. Marie, Ontario, Canada

[16]McGill University, Montréal, Québec, Canada

[17]Smithsonian Tropical Research Institute, Panama

[18]Department of Forest Resources, University of Minnesota, St. Paul, MN 55108, USA

[19] Hawkesbury Institute for the Environment, Western Sydney University, Penrith, NSW 2753, Australia

[20]Institute for Global Change Biology, and School for the Environment and Sustainability, University of Michigan, Ann Arbor, MI 48109, United States

**Contact information**

### Methods

We used funnel plots (*funnel* function in the package *metafor*) to assess the distribution of effect sizes in each of our models and considered the impact of removing the most extreme outliers on the results.

### Results

#### Research question 1a – mixed vs. average of monocultures

All comparisons included:

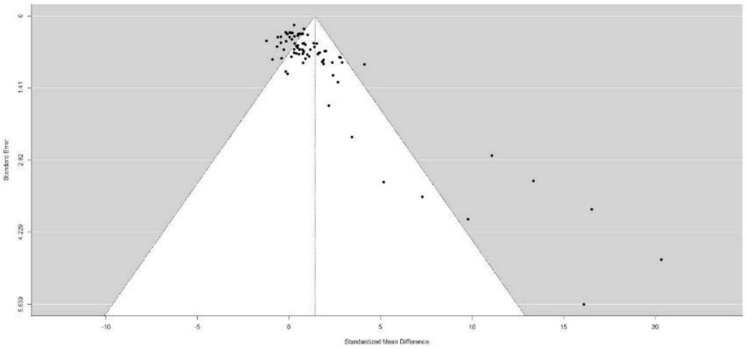


SMD 1.41 [95% CI 0.75, 2.07], k = 79

Removal of most extreme outliers, standardised mean difference > 15:

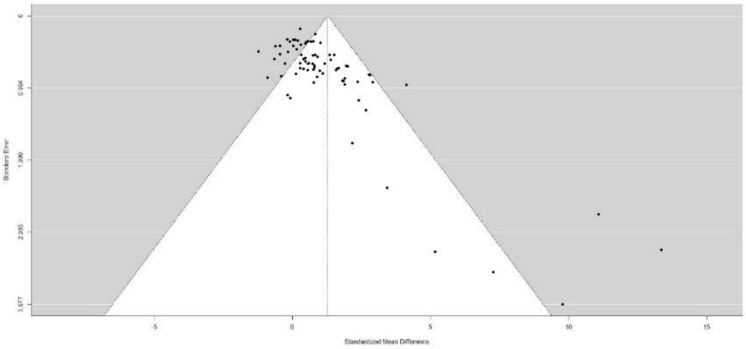


SMD 1.28 [95% CI 0.75, 1.82], k = 76

Removal of most extreme outliers, standardised mean difference > 10:

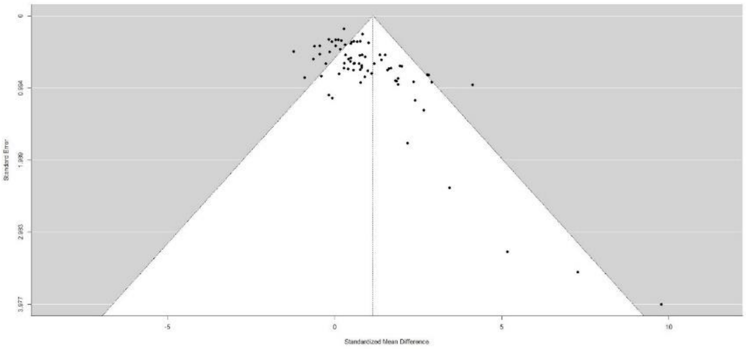


SMD 1.14 [95% CI 0.71, 1.56], k = 74

No qualitative change to results, carbon stocks in mixed plantation still clearly higher than the average of monocultures.

**Research question 1b – mixed vs. best monoculture**

All comparisons included:

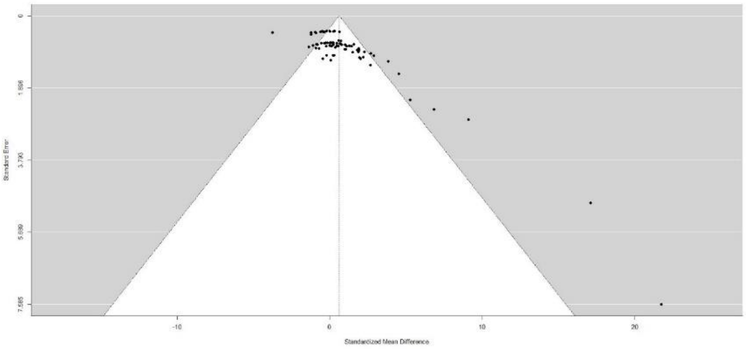


SMD 0.62 [95% CI −0.0095, 1.25], k = 79.

Removal of most extreme outliers, standardised mean difference > 10:

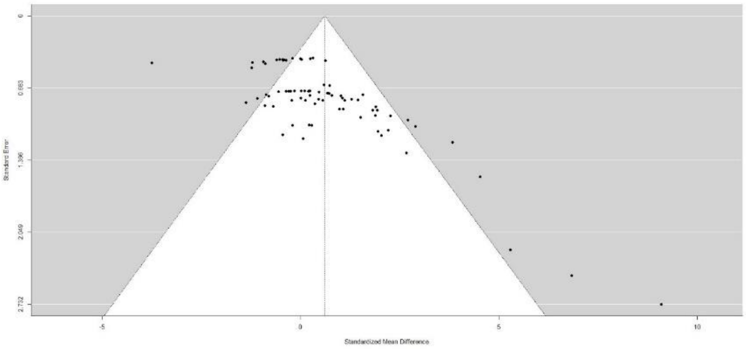


Pooled SMD 0.61 [95% CI −0.015, 1.23], k = 77.

Removal of additional outliers, standardised mean difference > 6 and < −3:

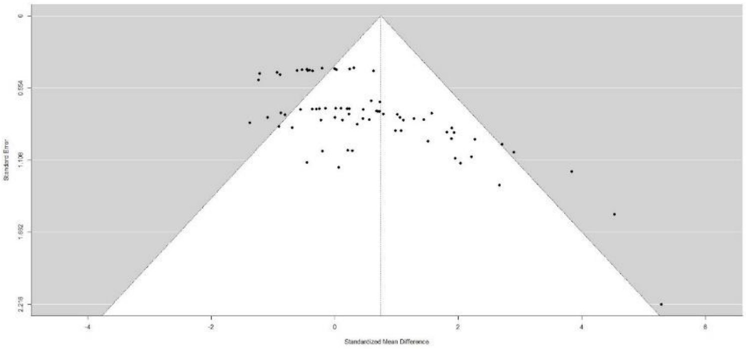


Pooled SMD 0.75 [95% CI 0.32, 1.17], k = 73.

Removal of these outliers alters the result to show higher carbon in the mixed plantation relative to best monoculture, with the confidence interval no longer overlapping zero. This contrasts to the result with the full dataset, which suggested no clear increase in carbon stocks in the mixed plantation relative to best monoculture.

**Research question 1c – mixed vs commercial monocultures**

All comparisons included:

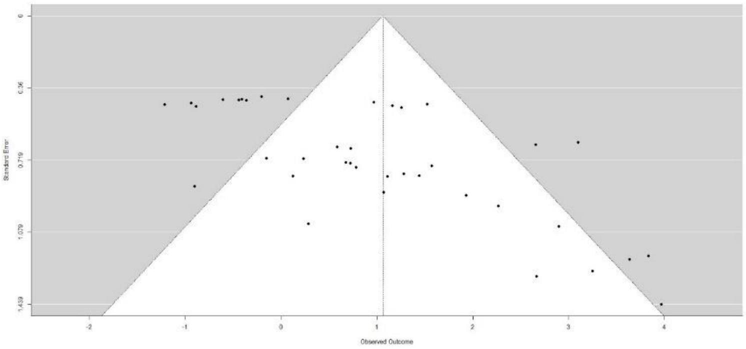


SMD 1.06 [0.47, 1.65], k = 38

Remove effect sizes < −0.5:

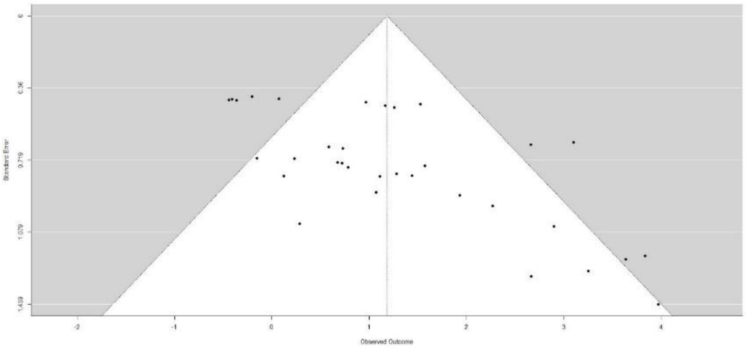


SMD 1.18 [95% CI 0.70, 1.67], k = 33

No qualitative change in results, still shows higher carbon stocks in mixed plantation relative to commercial monocultures.

**Research question 2 – effect of richness in mixture on carbons stocks**

*Effect of richness, mixed vs. average of monocultures*

All comparisons included:

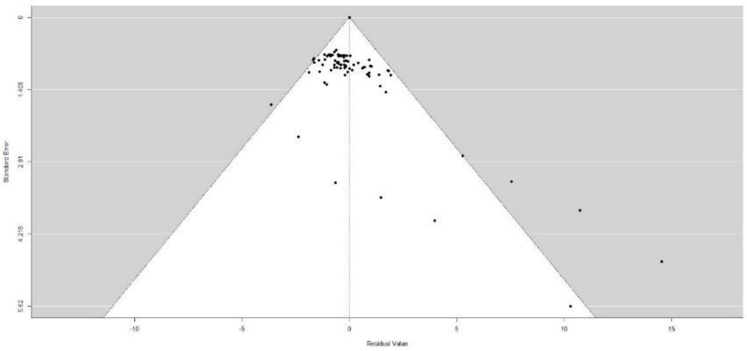

$Q_M = 46.79$, $p < 0.01$, $k = 79$

Remove effect sizes > 16.5

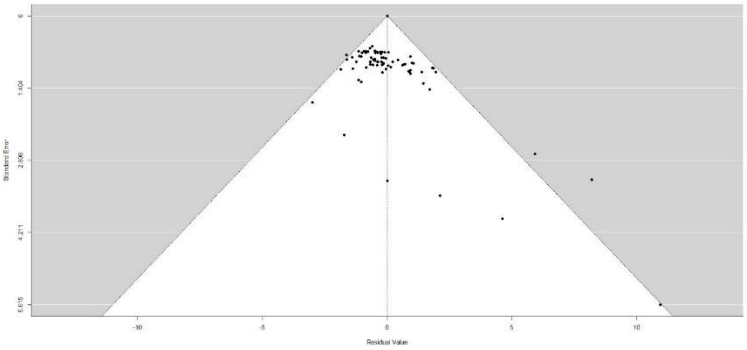

$Q_M = 39.22$, $p < 0.001$, $k = 77$

No qualitative change in results, still significant relationship between richness and carbon stocks in mixed vs. average of monocultures.

*Species richness 6, mixed vs. average of monocultures*

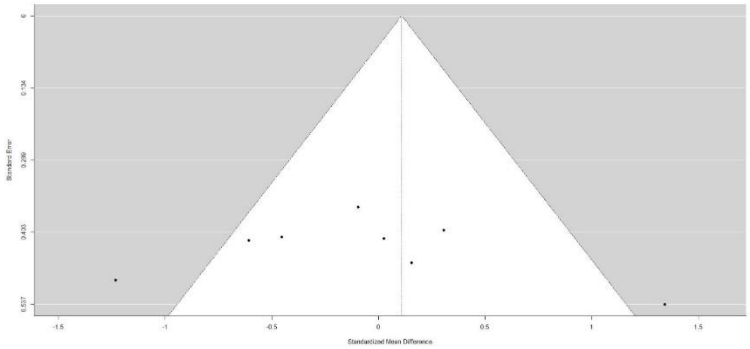


SMD 0.11 [95% CI −0.55, 0.77], k = 8

*Species richness 4, mixed vs. average of monocultures*

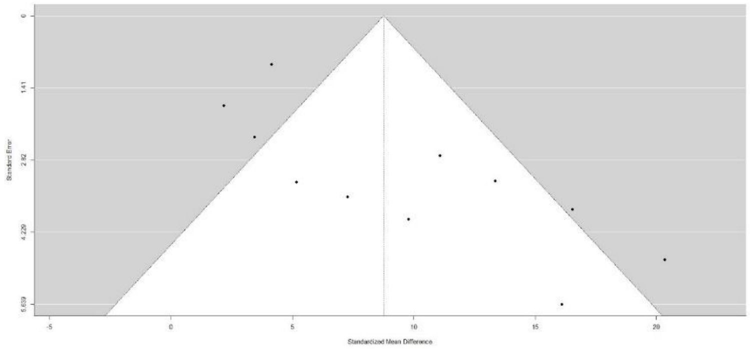


SMD 8.76 [95% CI 4.82, 12.7], k = 11

*Species richness 2, mixed vs. average of monocultures*

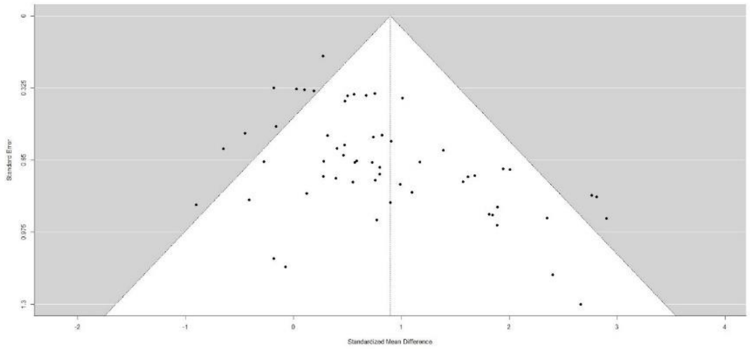


SMD 0.90 [0.53, 1.27], k = 58

*Effect of richness, mixed vs. best monoculture*

All comparisons included:

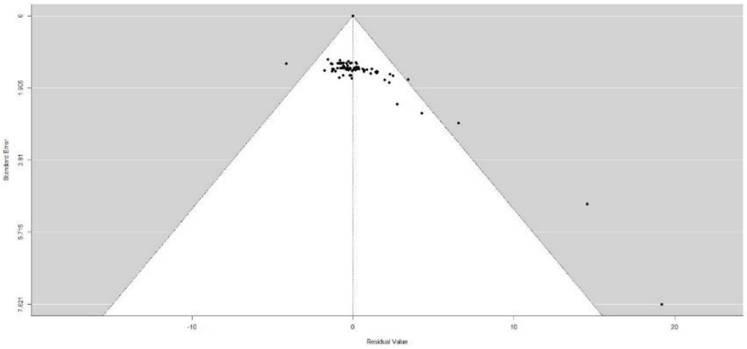


$Q_M$ 8.78, p = 0.067, k = 79

Remove effect sizes > 10:

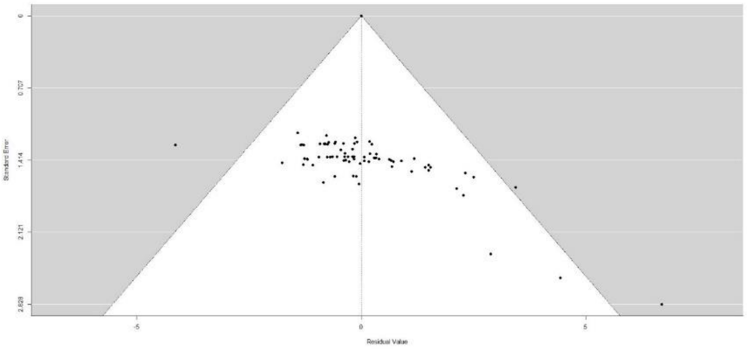


$Q_M$ = 8.25, p = 0.083, k = 77

Remove effect sizes > 6 and < −3:

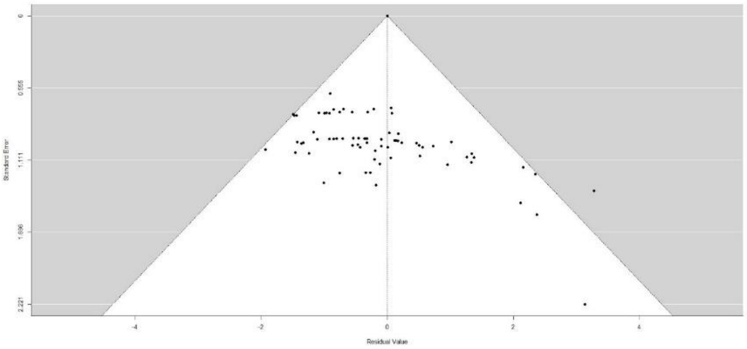


$Q_M$ = 11.25, p = 0.024, k = 74

Removal of these most extreme outliers alters the result, showing a significant relationship between richness and carbon stocks in mixed vs. best monoculture.

*Species richness 6, mixed vs. best monoculture*

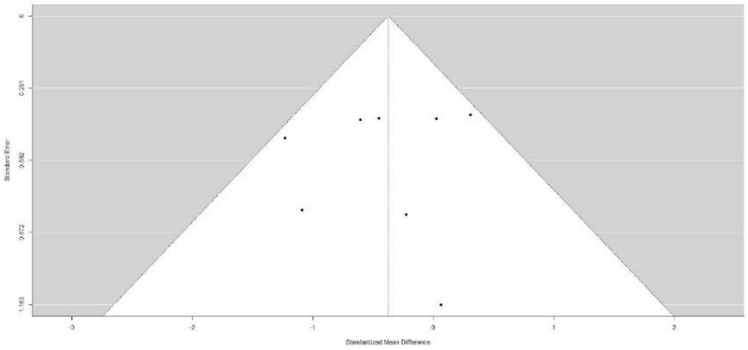

SMD −0.38 [95% CI −0.79, 0.042], k = 8

*Species richness 4, mixed vs. best monoculture*

All comparisons included

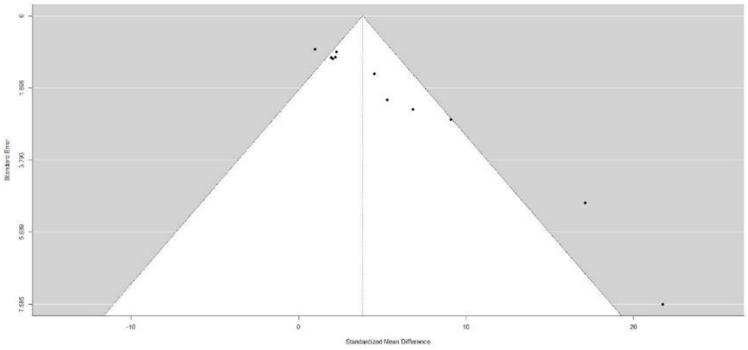

SMD 3.82 [95% CI 1.69, 5.95], k = 11

Remove effect sizes > 10:

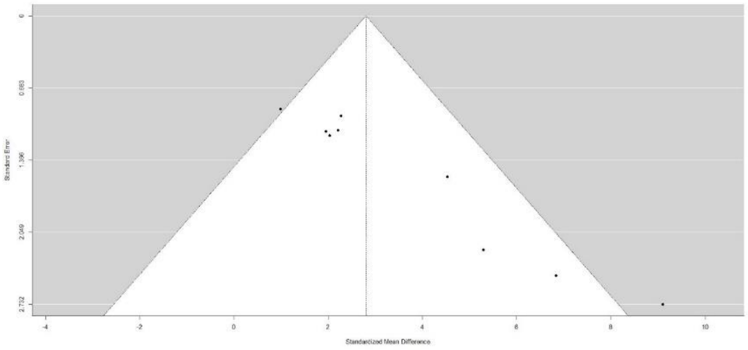

SMD 2.80 [95% CI 1.69, 3.91], k = 9

No qualitative change in results, still significantly higher carbon stocks in mixture relative to best monoculture for species richness 4.

*Species richness 2, mixed vs. best monoculture*

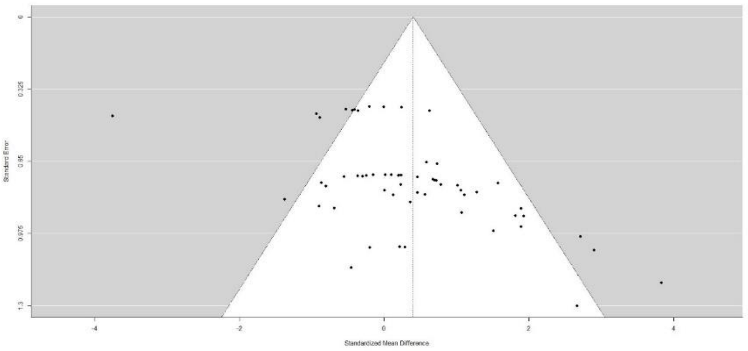


SMD 0.40 [95% CI −0.28, 1.08], k = 58

Remove effect sizes < −2:

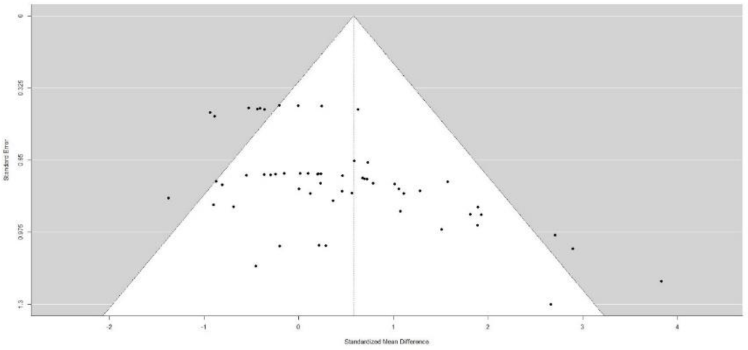


SMD 0.58 [95% CI 0.17, 0.99], k = 57

Removal of these most extreme outliers alters the result, showing greater carbon in mixed vs. best monoculture for species richness 2.

*Species richness 2, mixed vs. commercial monoculture*

All comparisons included:

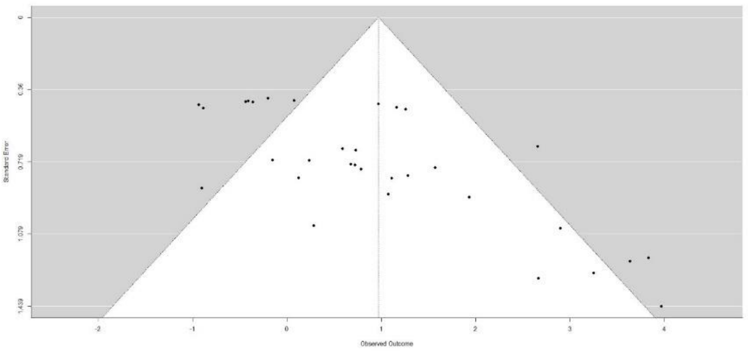


SMD 0.97 [95% CI 0.34, 1.61], k = 32

Remove effect sizes < −0.5

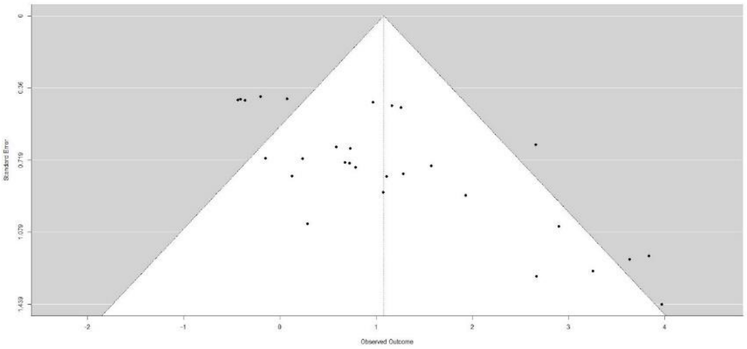


SMD 1.08 [95% CI 0.52, 1.64], k = 29

No qualitative change in results, still higher carbon stocks in mixed vs. commercial monocultures for species richness 2.

**Research question 3 – potential mechanisms**

*Inclusion of N-fixer in mixture, mixed vs. average of monocultures*

All comparisons included:

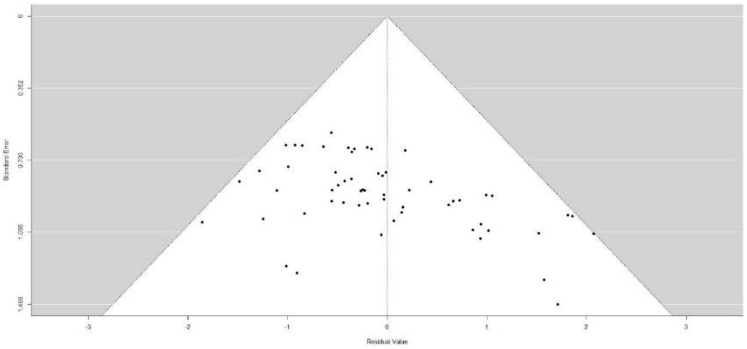


$Q_M$ = 0.37, p = 0.54, k = 58

*N-fixer present, mixed vs. average of monocultures*

All comparisons included:

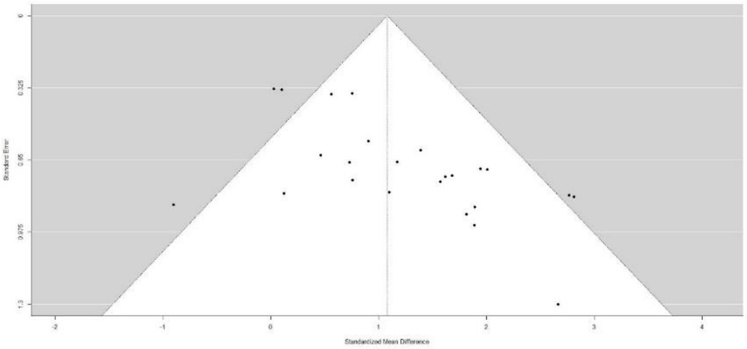


SMD 1.08 [95% CI 0.58, 1.58], k = 24

*N-fixer not present, mixed vs. average of monocultures*

All comparisons included:

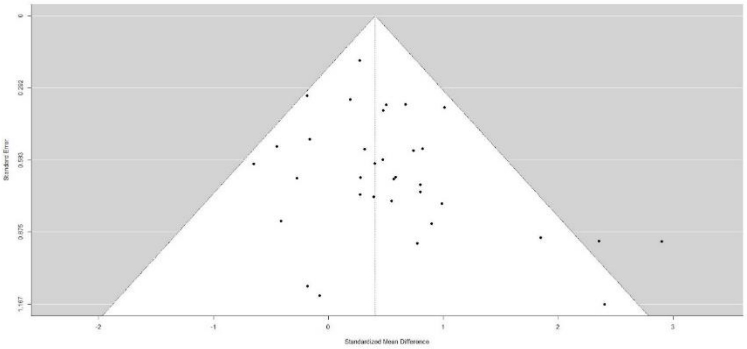

SMD 0.41 [95% CI 0.23, 0.58], k = 34

*Inclusion of N-fixer in mixture, mixed vs. best monoculture*

All comparisons included:

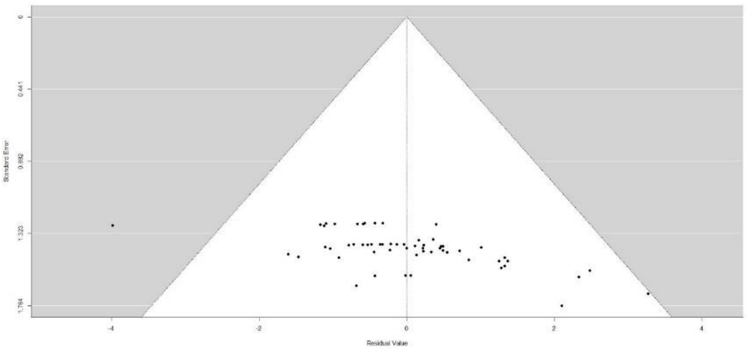

$Q_M$ = 1.35, p = 0.25, k = 58

Remove effect size < −3:

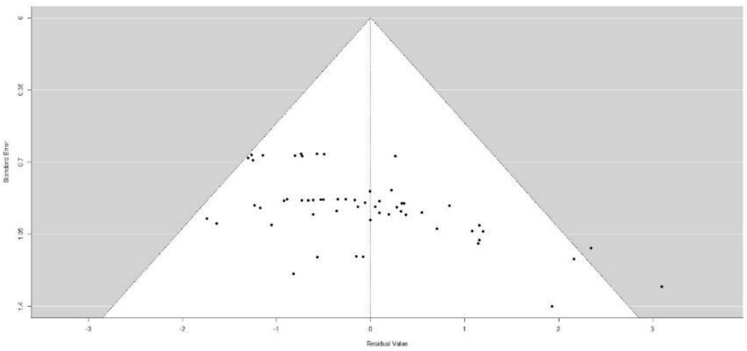

$Q_M$ = 2.25, p = 0.13, k = 57

No qualitative change in results, moderator for nitrogen-fixer present vs. absent still not significant for comparison between mixed and best monocultures.

*N-fixer present, mixed vs. best monoculture*

All comparisons included:

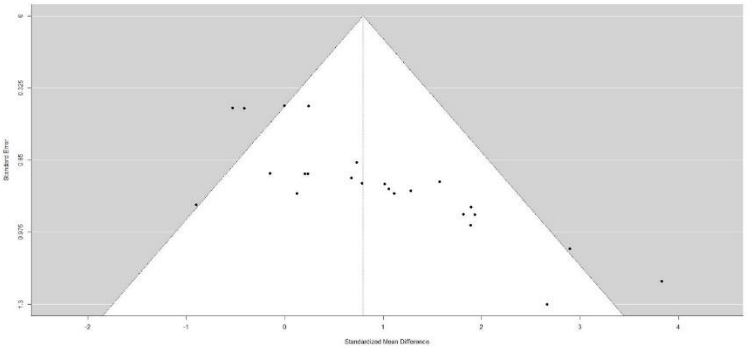


SMD 0.80 [95% CI 0.24, 1.36], k = 24

*N-fixer not present, mixed vs. best monoculture*

All comparisons included:

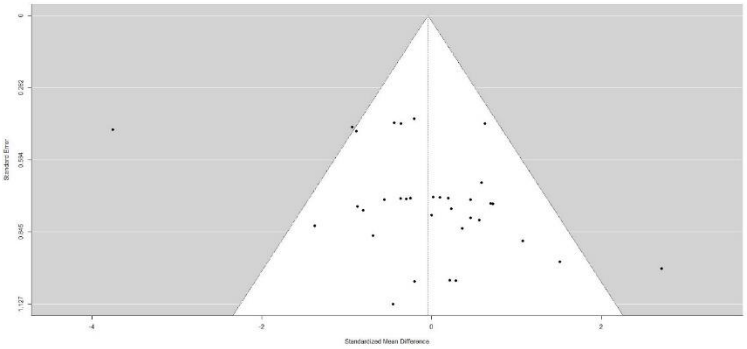


SMD −0.043 [95% CI −1.09, 1.00], k = 34

Remove effect size < −3:

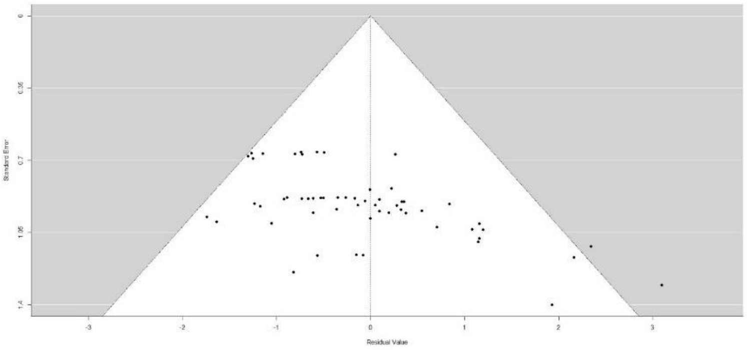


SMD 0.0098 [95% CI −0.28, 0.30], k = 33

No qualitative change in results, carbon stocks similar in mixed vs. best monoculture when N-fixer not present in the mixture.

*Inclusion of N-fixer in mixture, mixed vs. commercial monoculture*

All comparisons included:

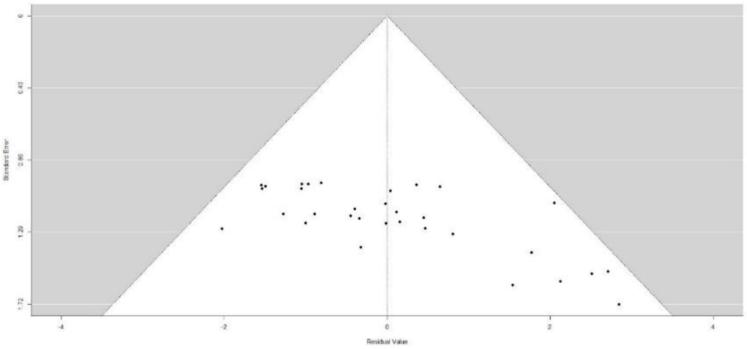

$Q_M = 1.33$, $p = 0.25$, $k = 32$

*N-fixer present, mixed vs. commercial monoculture*

All comparisons included:

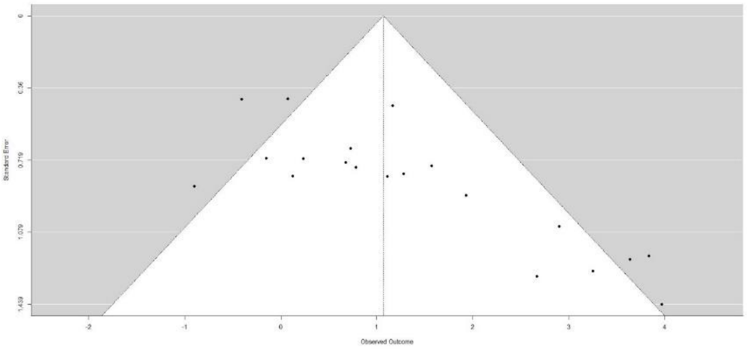

SMD 1.07 [95% CI 0.29, 1.84], k = 20

*N-fixer not present, mixed vs. commercial monoculture*

All comparisons included:

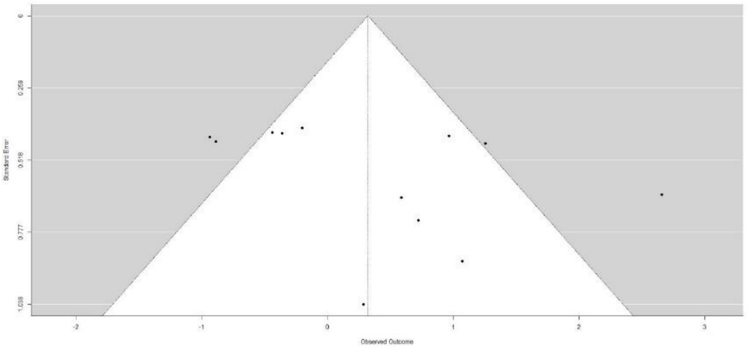

SMD 0.32 [95% CI −0.28, 0.92], k = 12

Remove effect size > 2:

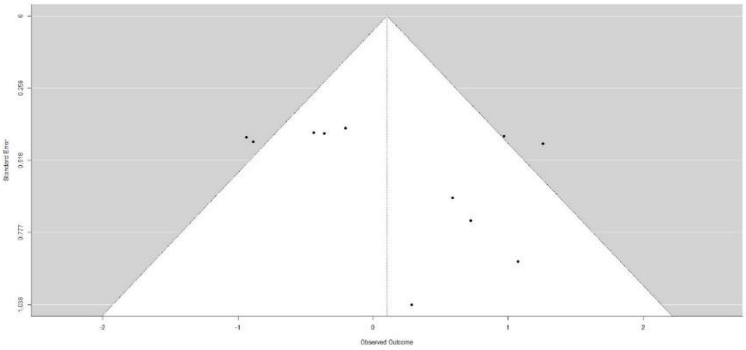

SMD 0.10 [95% CI −0.42, 0.63], k = 11

No qualitative change in results, carbon stocks similar in mixed vs. commercial monocultures when N-fixer not present in the mixture.

*Effect of species origin in mixture, mixed vs. average of monocultures*

All comparisons included:

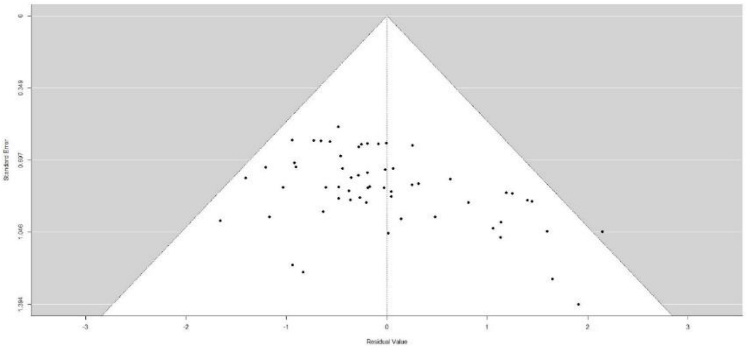

$Q_M = 0.94$, $p = 0.16$, $k = 58$

*Native only in mixture vs. average of monocultures*

All comparisons included:

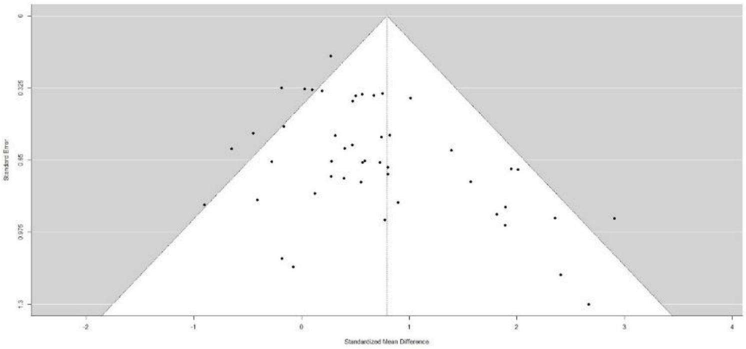

SMD 0.79 [95% CI 0.36, 1.22], k = 47

*Non-native/mixed origin in mixture vs. average of monocultures*

All comparisons included:

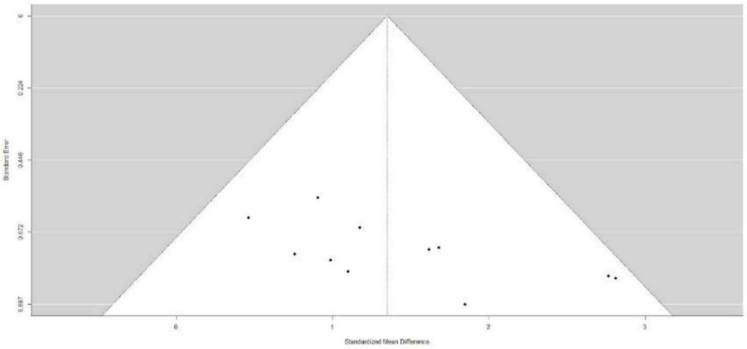

SMD 1.35 [95% CI 0.92, 1.78], k = 11

*Species origin in mixture, mixed vs. best monoculture*

All comparisons included:

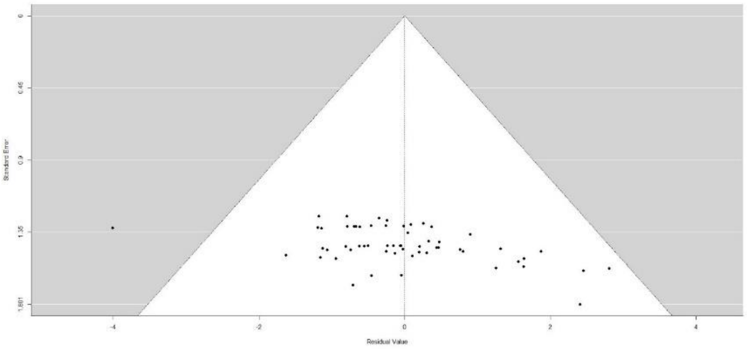

$Q_M = 0.75$, $p = 0.39$, $k = 58$

Remove effect size < −3:

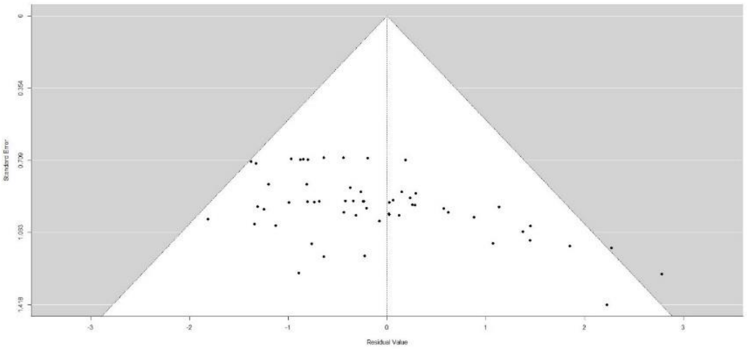

$Q_M = 1.54$, $p = 0.21$, $k = 57$

No qualitative change in results, moderator for species origin still not significant for comparison between mixed and best monocultures.

*Native only in mixture vs. best monoculture*

All comparisons:

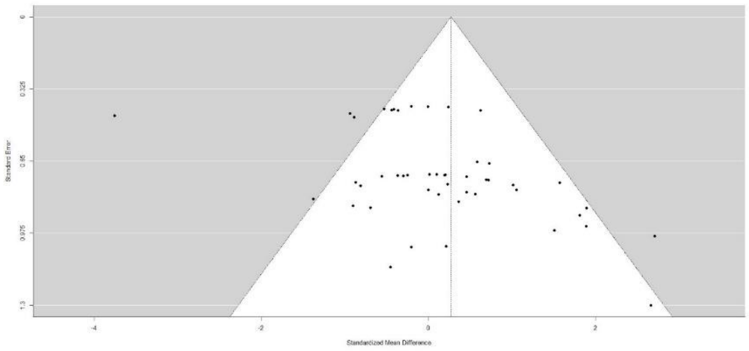

SMD 0.27 [95% CI  -0.53, 1.08], k = 47

Remove effect size < −3:

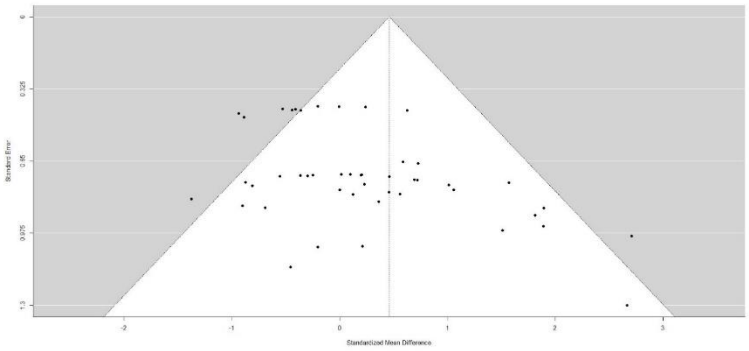

SMD 0.46 [95% CI 0.0008, 0.92], k = 46

No qualitative change in results, carbon stocks similar in mixed vs. best monoculture when all native species in mixture.

*Non-native/mixed origin in mixture vs. best monoculture*

All comparisons included:

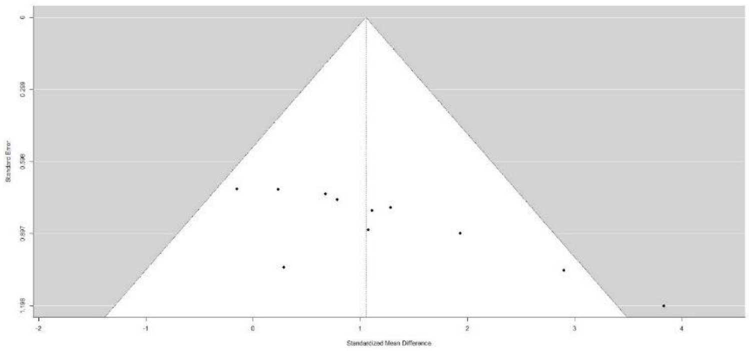

SMD 1.05 [95% CI 0.45, 1.65], k = 11

*Species origin in mixture, mixed vs. commercial monocultures*

All comparisons included:

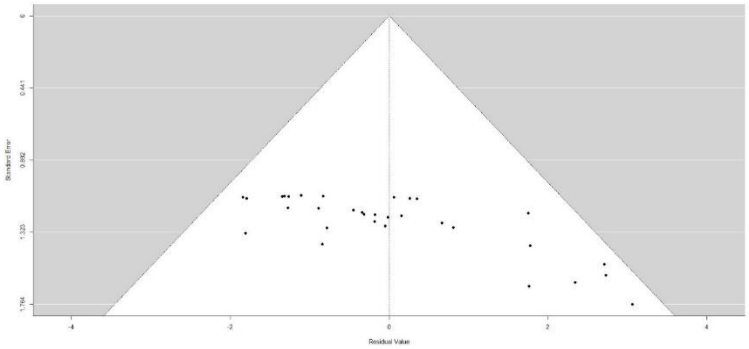


$Q_M = 0.091$, $p = 0.76$, $k = 32$

*Native only in mixture, mixed vs. commercial monocultures*

All comparisons included:

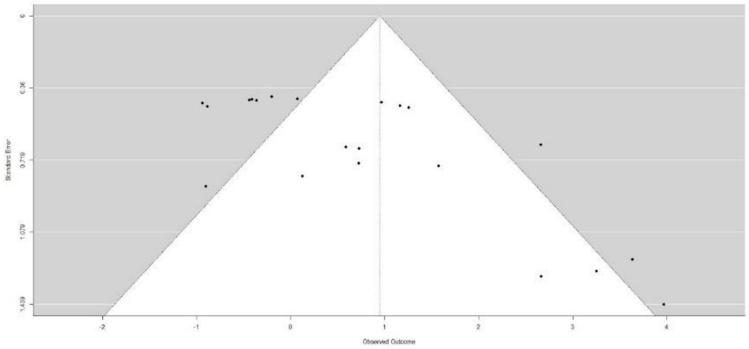


SMD 0.95 [95% CI −0.0072, 1.90], k = 21

Remove effect sizes < −0.5:

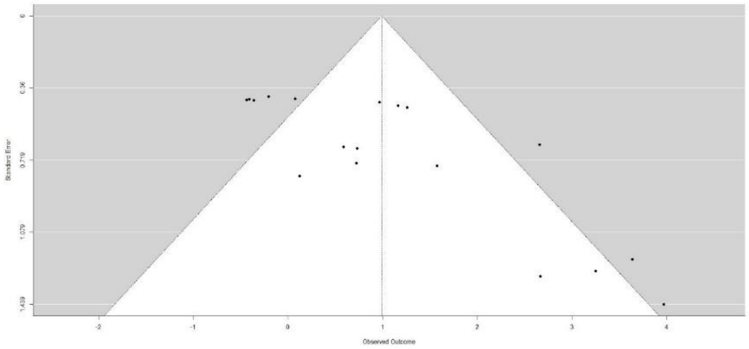


SMD 0.99 [95% CI 0.26, 1.71], k = 18

Remove effect sizes < 0:

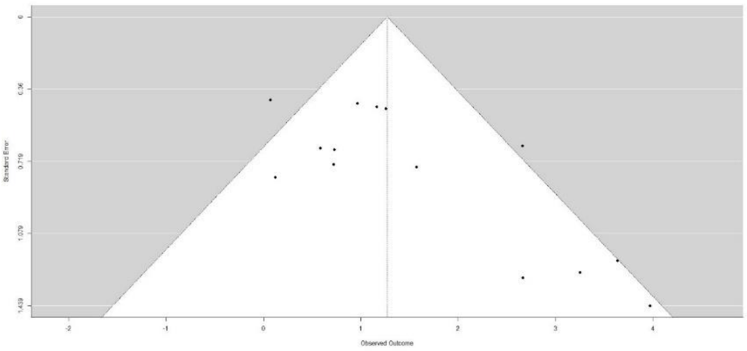

SMD 1.27 [95% CI 0.61, 1.94], k = 14

Removal of these outliers alters the result to show higher carbon in the mixed plantation relative to commercial monoculture when considering native species only in the mixture, with the confidence interval no longer overlapping zero. This contrasts to the result with the full dataset, which suggested no clear increase in carbon stocks in the mixed plantation relative to monoculture.

*Non-native/mixed origin in mixture vs. commercial monocultures*

All comparisons:

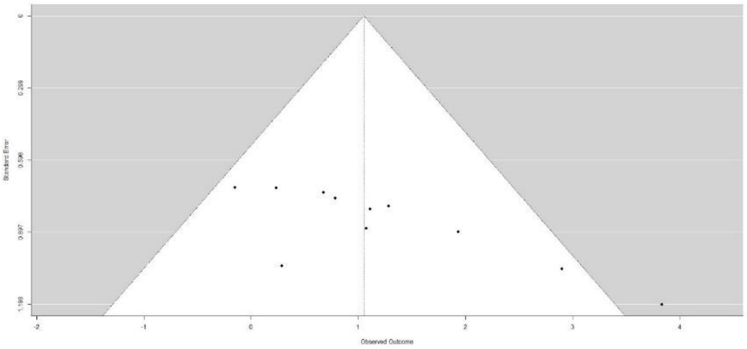

SMD 1.05 [95% CI 0.45, 1.65], k = 11

Remove effect size > 3:

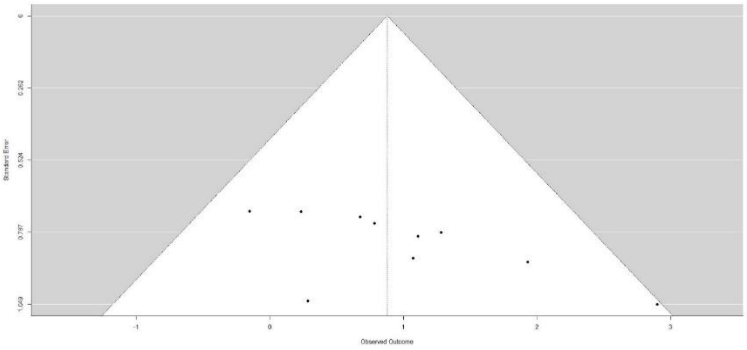

SMD 0.88 [95% CI 0.37, 1.38], k = 10

No qualitative change in results, carbon stocks similar in mixed vs. best monoculture when all non-native/mixed origin species in mixture.

*Effect of study design, mixed vs. average of monocultures*

All comparisons included:

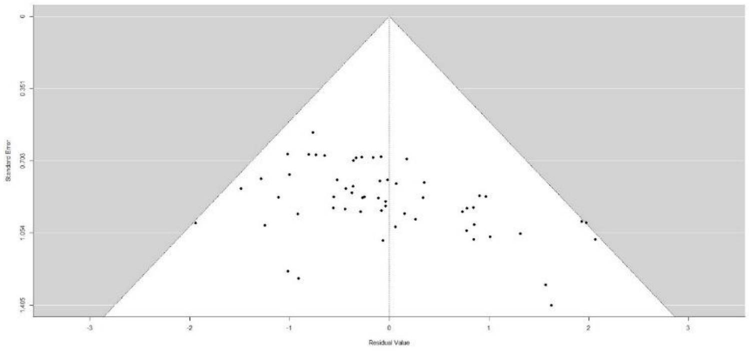

$Q_M = 0.24$, $p = 0.62$, $k = 58$

*Designed experiment only, mixed vs. average of monocultures*

All comparisons included:

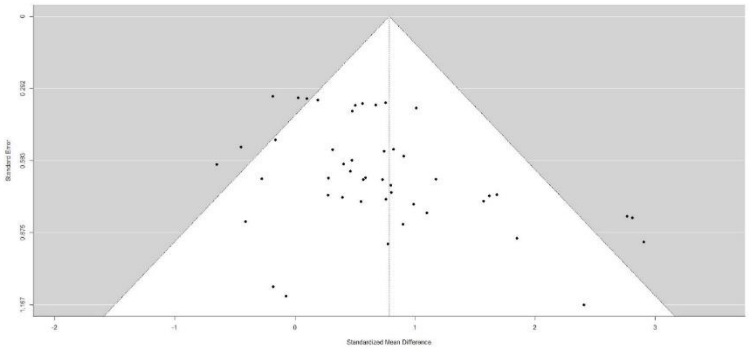

SMD 0.78 [95% CI 0.41, 1.16], k = 47

*Existing plantation only, mixed vs. average of monocultures*

All comparisons included:

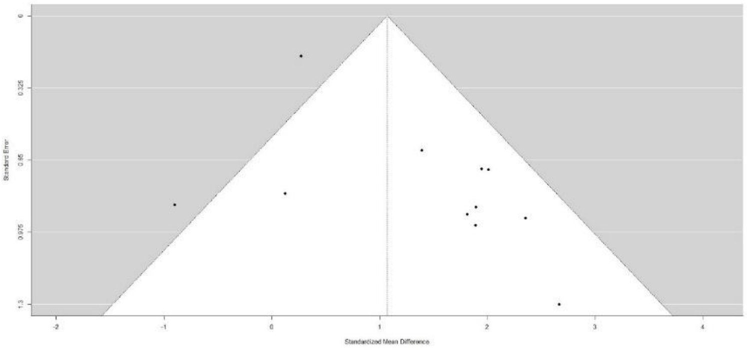


SMD 1.07 [95% CI 0.19, 1.95], k = 11

*Effect of study design, mixed vs. best monoculture*

All comparisons included:

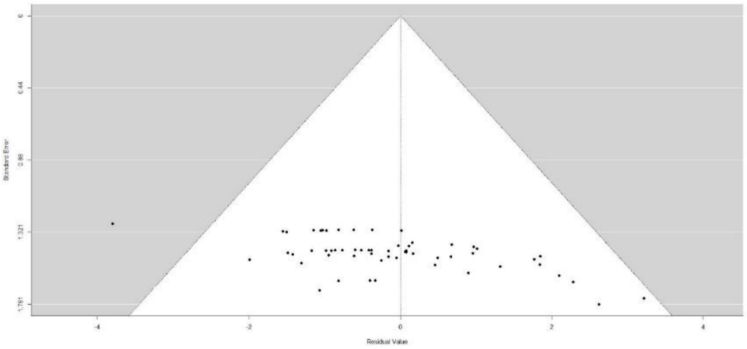


$Q_M$ = 0.63, p = 0.43, k = 58

Remove effect sizes < −3:

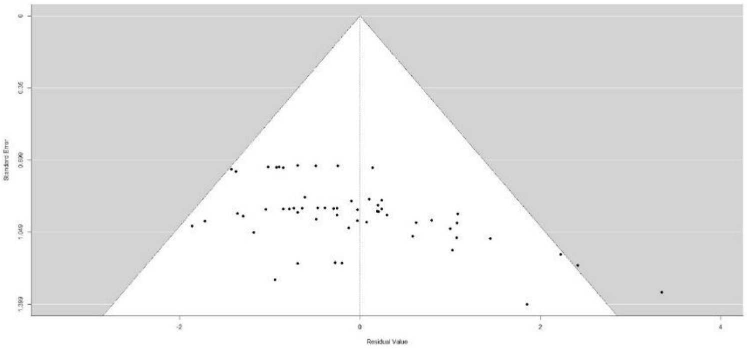


$Q_M$ = 0.51, p = 0.48, k = 57

Remove effect size > 3:

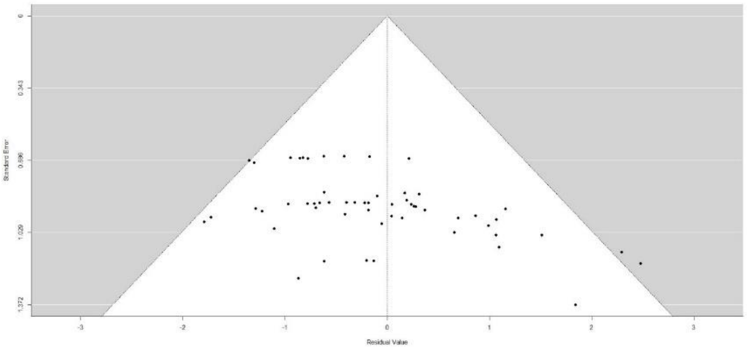


$Q_M = 0.88$, $p = 0.35$, $k = 56$

No qualitative change in results, moderator for study design still not significant for comparison between mixed and best monocultures.

*Designed experiment only, mixed vs. best monoculture*

All comparisons included:

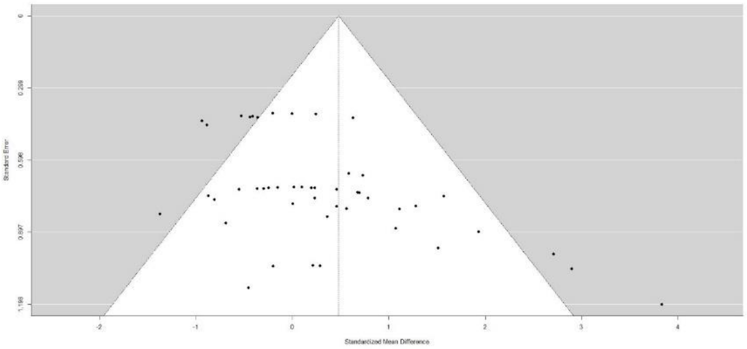


SMD 0.48 [95% CI 0.013, 0.95], k = 47

Remove effect size > 3:

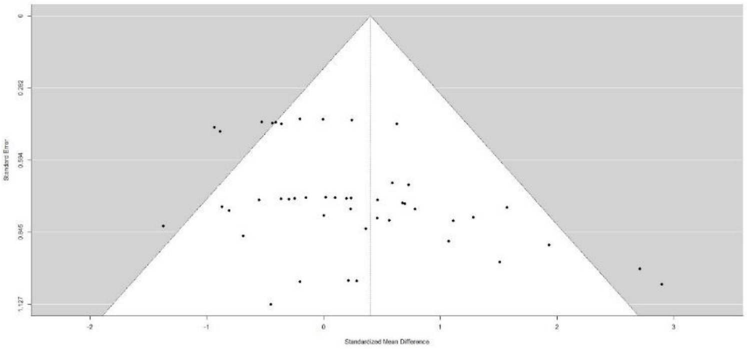


SMD 0.40 [95% CI −0.016, 0.82], k = 46

Removal of outlier alters the result to show no clear difference in carbon in the mixed plantation relative to best monoculture when considering designed experiments only, with the confidence interval overlapping zero. This contrasts to the result with the full dataset, which suggested higher carbon stocks in the mixed plantation relative to monoculture.

*Existing plantation only, mixed vs. best monoculture*

All comparisons included:

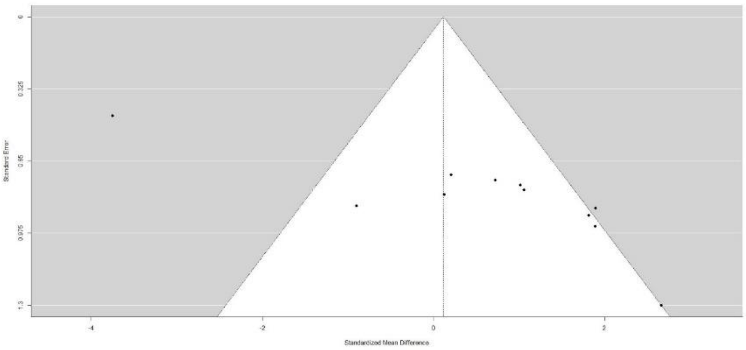


SMD 0.12 [95% CI −1.46, 1.69], k = 11

Remove effect size < −3:

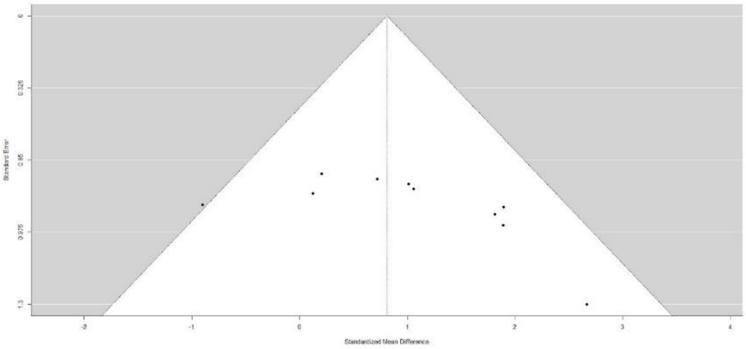


SMD 0.81 [95% CI −0.015, 1.64], k = 10

No qualitative change in results, carbon stocks similar in mixed vs. best monoculture for existing plantations.

*Effect of study design, mixed vs. commercial monocultures*

All comparisons included:

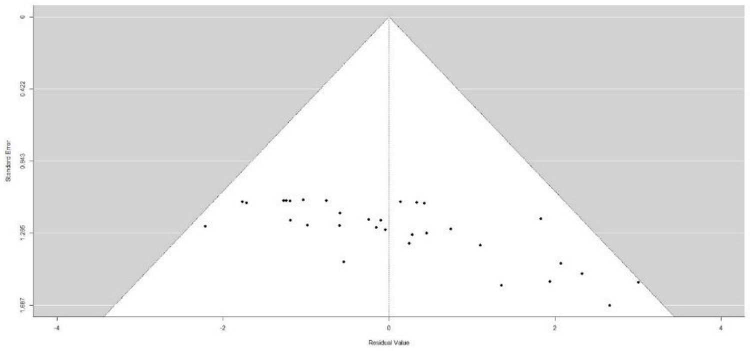


$Q_M = 0.46$, $p = 0.50$, $k = 32$

*Designed experiment only, mixed vs. commercial monocultures*

All comparisons included:

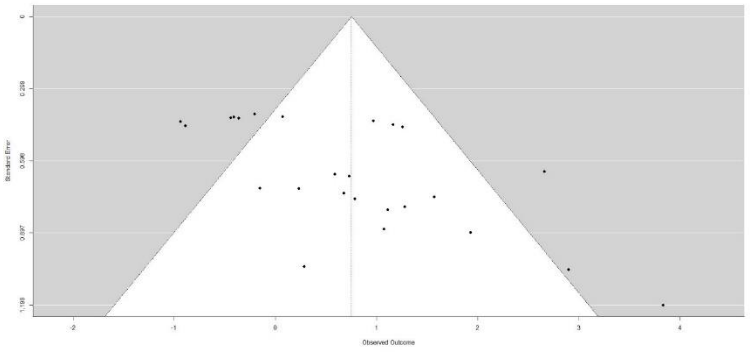


SMD 0.75 [95% CI 0.19, 1.31], k = 25

*Existing plantation only, mixed vs. commercial monocultures*

All comparisons included:

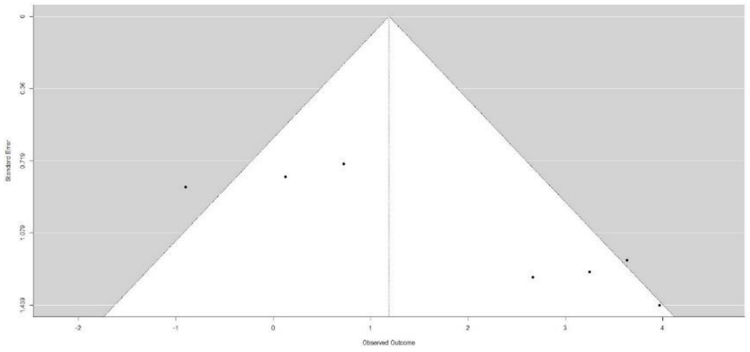


SMD 1.19 [95% CI −0.45, 2.83], k = 7#

*Effect of age, mixed vs. average of monocultures*

All comparisons included:

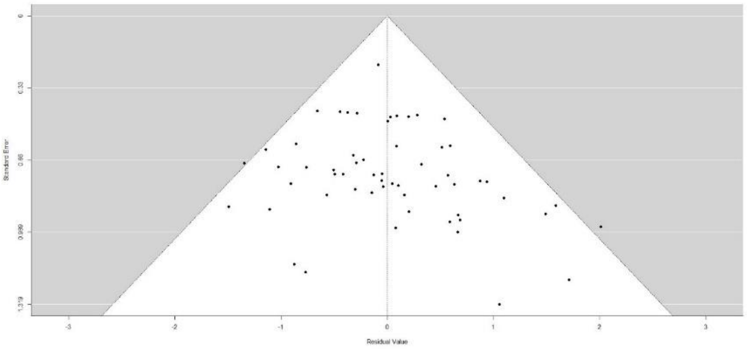


$Q_M$ = 13.3, p = 0.001, k = 58, quadratic term −0.013 [−0.023, −0.003]

*Effect of age, mixed vs. best monoculture*

All comparisons included:

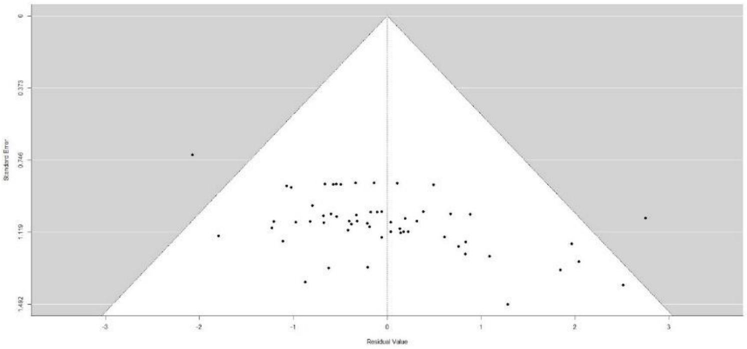


$Q_M$ = 16.1, p < 0.001, k = 58, quadratic term −0.019 [95% CI −0.029, −0.008]

Remove effect sizes > 3 and < −3:

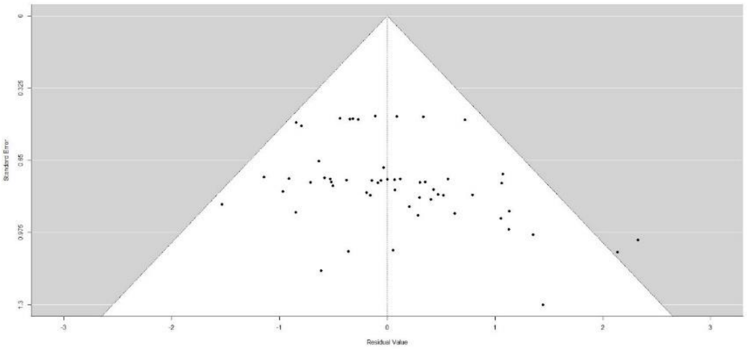


$Q_M$ = 22.1, p < 0.001, k = 56, quadratic term −0.012 [95% CI −0.019, −0.006]

No qualitative change in results, moderator for age still significant for comparison between mixed and best monoculture.

*Effect of age, mixed vs. commercial monocultures*

All comparisons included:

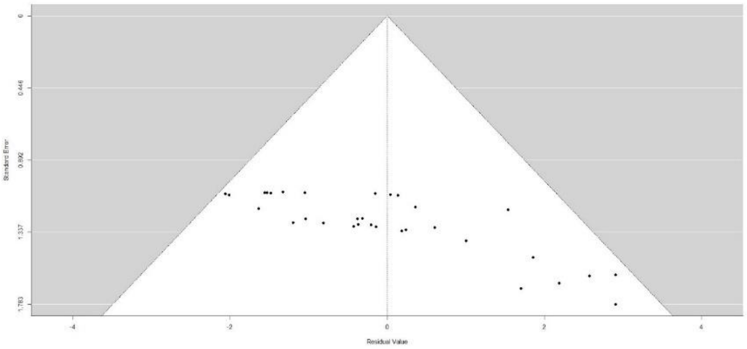


$Q_M = 0.14$, $p = 0.71$, $k = 32$, effect Age −0.02 [95% CI −0.12, 0.082]
